# Restoring mechanophenotype reverts malignant properties of ECM-enriched vocal fold cancer

**DOI:** 10.1101/2024.08.22.609159

**Authors:** Jasmin Kaivola, Karolina Punovuori, Megan R. Chastney, Yekaterina A. Miroshnikova, Hind Abdo, Fabien Bertillot, Fabian Krautgasser, Jasmin Di Franco, James R.W. Conway, Gautier Follain, Jaana Hagström, Antti Mäkitie, Heikki Irjala, Sami Ventelä, Hellyeh Hamidi, Giorgio Scita, Roberto Cerbino, Sara A. Wickström, Johanna Ivaska

## Abstract

Increased extracellular matrix (ECM) and matrix stiffness promote solid tumor progression. However, mechanotransduction in cancers arising in mechanically active tissues remains underexplored. Here, we report upregulation of multiple ECM components accompanied by tissue stiffening in vocal fold cancer (VFC). We compare non-cancerous (NC) and patient-derived VFC cells – from early (mobile, T1) to advanced-stage (immobile, T3) cancers – revealing an association between VFC progression and cell-surface receptor heterogeneity, reduced laminin-binding integrin cell-cell junction localization and a flocking mode of collective cell motility. Mimicking physiological movement of healthy vocal fold tissue (stretching/vibration), decreases oncogenic nuclear β-catenin and YAP levels in VFC. Multiplex immunohistochemistry of VFC tumors uncovered a correlation between ECM content, nuclear YAP and patient survival, concordant with VFC sensitivity to YAP-TEAD inhibitors in vitro. Our findings present evidence that VFC is a mechanically sensitive malignancy and restoration of tumor mechanophenotype or YAP/TAZ targeting, represents a tractable anti-oncogenic therapeutic avenue for VFC.

---

The human vocal folds are composed of three layers (epithelial layer, basement membrane and lamina propria) with distinct cellular and extracellular matrix (ECM) compositions^1^. Maintaining proper ECM organization is essential for vocal fold epithelium viscoelasticity, as it has been shown that the biomechanical and physiological performance of the vocal folds relies on ECM homeostasis^2,3^. ECM alterations are also linked to numerous pathological conditions, such as cancer^4^. Vocal fold cancer (VFC) remains a major clinical challenge with limited targeted therapy options, and only a 34% 5-year survival rate for advanced T3-T4 disease. VFC arises in the stratified squamous epithelium, and as it progresses, the squamous cells in the epithelial layer breach the underlying basement membrane, invade into the collagen-rich lamina propria and further to the underlaying muscle, leading to mechanical fixation^4–7^, characteristic to T3 and T4 disease.

In recent years, there has been a growing appreciation of the role of ECM remodeling and increased ECM deposition in cancer pathogenesis^8,9^ as the ensuing increase in tissue rigidity alters tissue mechanics and drives cancer progression^10–13^. Integrins, the main cellular ECM receptors^14^, act as mechanosensors by probing the physical properties of their surroundings and transducing this information via the cytoskeleton into intracellular biochemical signals and transcriptional changes^15–17^. Among the key oncogenic signals triggered by increased tissue rigidity and integrin engagement, is stabilization and nuclear translocation of the hippo-signaling pathway transcription factors YAP and TAZ^18,19^. YAP/TAZ are upregulated in various cancers and influences tumor initiation, progression and therapeutic resistance^20–22^. Importantly, this signaling is reciprocal with YAP positive control of focal adhesion (FA) assembly^23^ and integrin adhesion to the ECM regulates YAP/TAZ in the squamous epithelium^24^. However, it remains unknown whether changes in ECM and cell mechanics play a role in VFC. Further, it is not known whether immobility caused by fixation contributes to VFC malignancy or correlates with patient outcome.

The role of ECM and mechanical forces in tumor development, have predominantly been investigated in solid tumors arising from non-motile tissues such as the mammary gland, brain, and pancreas with a focus solely on the outcomes of increased rigidity. Recently, continuous dynamic mechanical challenge to the lung epithelium was shown to increase nuclear YAP in ventilated rat lungs^25^ and cell stretching was shown to trigger changes in heterochromatin architecture and nuclear softening^26^. In contrast, the role of mechanical stimuli on cancer progression in mechanically active organs, which are under continuous biomechanical stress, has not been explored. Due to the unique biomechanical properties of the vocal fold, we sought to understand the role of cell-matrix- and cell-cell adhesion and their mechanical regulation in VFC. We predicted that vocal fold epithelial cell responses to dynamic mechanical vibration and stretching, akin to the situation in vivo, would deviate significantly from currently established principles of cell mechanobiology. Moreover, we set out to explore whether mechanical stimuli would be essential not only for phonation but for tissue homeostasis and whether restoration of mechanical stress in advanced mechanically fixed VFC, would reverse the oncogenic properties of these cells.

## Results

### Vocal fold cancer is associated with elevated gene expression of ECM components and stiffening of tissue

Earlier studies have demonstrated that vocal fold trauma, such as scarring, can lead to fibronectin and collagen accumulation in the tissue^3,27^. Moreover, VFC progression causes vocal fold immobility as the squamous cell carcinoma invades the underlying muscle and tissues of the neck. VFC staging is based on the mobility status of the vocal folds and invasion of surrounding tissues; in T1-T2 the vocal folds move normally, whereas in T3-T4 mechanical fixation renders the vocal fold(s) immobile (**Fig.1a & b**). We aimed to investigate the ECM composition and stiffness of VFC tissue compared to normal tissue in patient samples. First, we analyzed head and neck cancer RNA-sequencing data generated by The Cancer Genome Atlas (TCGA) research^28^, focusing specifically on samples with patient reports mentioning involvement of the vocal fold tissue (glottic larynx). Considering the low number of T1 and T2 cancer samples (n=4), we pooled all cancer samples together. Normal (n=12) and cancer (T1-T4, n=54) samples were compared to determine differentially expressed genes; 2041 genes were upregulated and 1629 downregulated in cancer samples compared to normal tissue (false discovery rate, FDR < 0.05). Gene ontology (GO) enrichment analysis^29,30^ revealed ECM and collagen-related GO-terms such as collagen-containing extracellular matrix, basement membrane and protein complex involved in cell adhesion, over-represented in the upregulated genes in cancer (**Fig.1c**). Conversely, over-represented GO-terms in the downregulated genes were linked to cell junctions and apical regions of the cell (**Fig.1d**). We further determined the genes encoding ECM and ECM-associated proteins in the data set using Matrisome AnalyzeR^31,32^. Strikingly, all differentially expressed collagens were upregulated including collagens I, III, IV and V that are abundant in the vocal folds^33^ (**Fig.1e**). Among the 76 differentially expressed ECM glycoprotein genes, 53 were upregulated and 23 downregulated (**Fig.1f; Extended data Fig.1a**). The upregulated genes included fibronectin (FN) and laminin-332 chains (LAMA3, LAMB3 and LAMC2), which can function as autocrine tumor promoters in squamous cell carcinoma^34^ through laminin-binding integrins α6β4 and α3β1. Moreover, 59 ECM regulator genes were upregulated (**Fig.1g**) and 28 downregulated (**Extended data Fig.1b**). The upregulated lysyl oxidases (LOXs) (LOX, LOXL, LOX2, and LOXL3), which covalently crosslink collagens to elastin, and metalloproteinases (MMP14, MMP2, MMP10, MMP1, MMP7, MMP19, MMP9, MMP12, MMP11, MMP13, MMP3, MMP17, MMP16 and MMP8) collectively allude to extensive ECM remodeling and stiffening in the cancerous tissue compared to normal tissue.

To further investigate the changes in ECM composition on the cellular level, we compared T1 (UT-SCC-11; 58-year old male) and T3 (UT-SCC-103; 51-year old male) patient-derived VFC cell lines, generated at the University of Turku^35–37^, to non-cancerous (NC) (HaCaT) cells. Western blot analysis confirmed fibronectin upregulation in T3 cancer cells in comparison to NC cells and T1 cancer cells (**Extended data Fig.1c & d**). Several collagens were also upregulated in our RNA-sequencing analysis (**Extended data Fig.1e**). To investigate if the altered ECM production impacted tissue stiffness, we performed atomic force microscopy (AFM) on patient NC (n=3) and cancer (n=2) samples (obtained from vocal fold surgery). Measurements of the elastic modulus confirmed a 3.2- fold increase in stiffness in cancer tissue (2.441 ± 1.479 kPa) in comparison to normal tissue (0.751 ± 0.341 kPa) (**Fig. 1h & i**). Taken together, these results demonstrate ECM component over- expression and significant tissue stiffening in VFC.

**Fig. 1.**
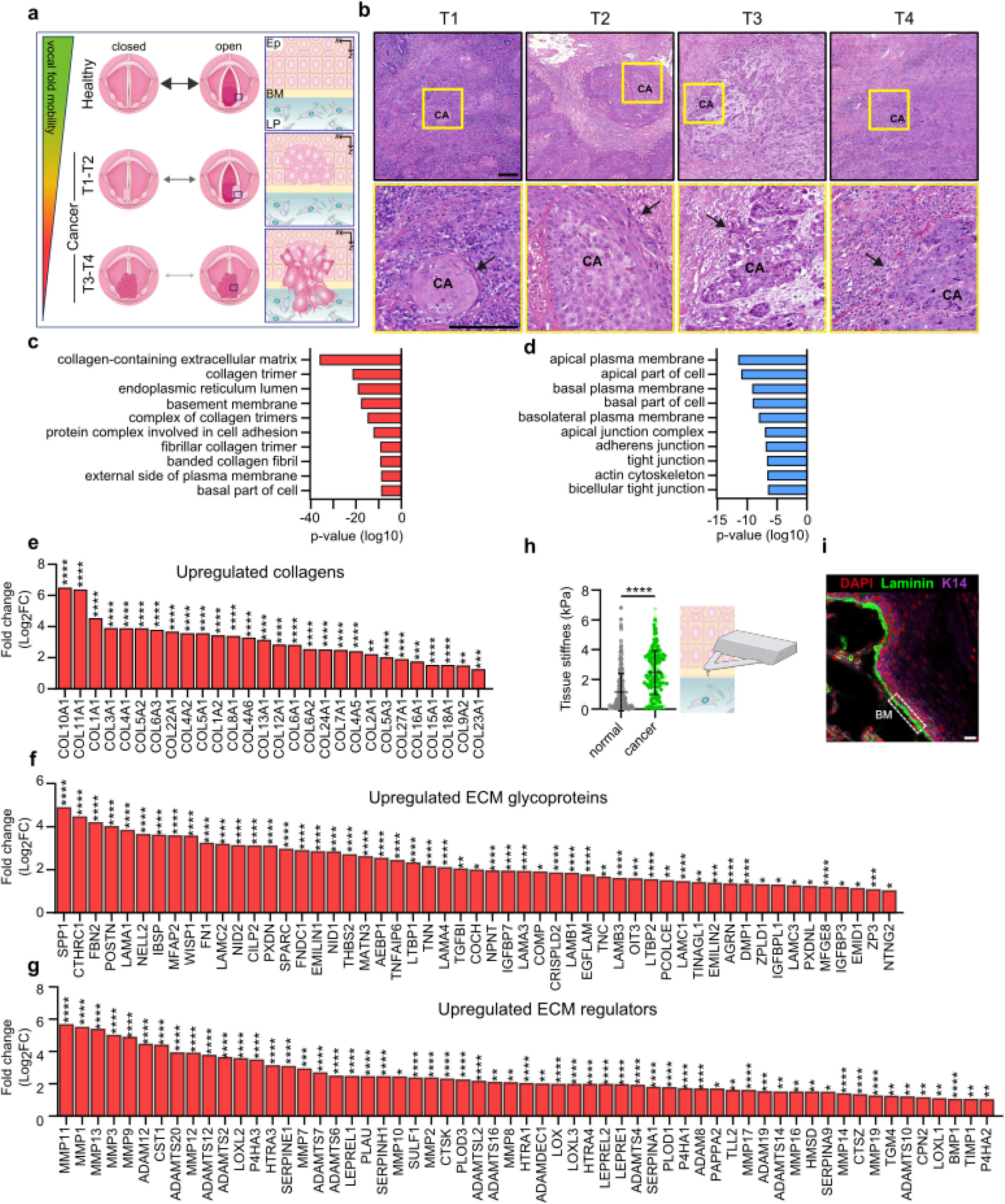
Vocal fold cancer is associated with elevated gene expression of ECM components and stiffening of tissue **a**, Schematic of changes in vocal fold mobility and invasion of transformed squamous cells through the basement membrane in VFC progression from T1 to T4. Ep= epithelium, BM= basement membrane, LP= lamina propria. **b**, Representative hematoxylin and eosin staining of T1-T4 vocal fold squamous cell carcinoma (CA) in patient tissue with arrows highlighting invasion. Scale bar 0.200 mm **c & d**, Over- represented GO-terms in upregulated (**c**) and downregulated (**d**) differentially expressed genes in VFC (T1- T4, n=54) compared to normal (n=12) patient tissue (TCGA-data, FDR < 0.001). **e-g**, Differentially upregulated (fold change, log2) collagens (**e**), ECM glycoproteins (**f**) and ECM regulators (**g**) in VFC (T1-T4, n=54) compared to normal (n=12) patient tissue (TCGA-data, FDR < 0.05) annotated with Matrisome analyzer^31^. **h**, Tissue stiffness (Pa) of normal (n=3) and cancer (n=2) vocal fold patient tissue measured by AFM. **i**, Representative immunofluorescence staining (dapi, laminin, K14) of normal vocal fold tissue. Scale bar 30 µm. Data are mean (± s.d.). FDR was used for assessment of statistical significance for differentially expressed genes and Mann-Whitney U-test for AFM measurements.

### Expression and subcellular localization of laminin-binding integrins is altered in vocal fold cancer

Guided by the differentially regulated genes identified in the TCGA-data associated with cell adhesion (**Fig. 1c**), we set out to explore the role of integrin adhesion complexes (IACs) in VFC. The patient data indicated upregulation of several genes encoding integrin adhesome proteins^38,39^, including an increase in laminin-binding integrins α3, α6 and β4. Integrin-α6β4 heterodimer is found in hemidesmosomes whereas integrins α3 and α6 form dimers with integrin β1in focal contacts^40,41^ (**Fig. 2a & b**). To determine whether these changes were recapitulated in the patient-derived cell lines, we used mass cytometry for high-dimensional phenotypic analysis of the cell-surface expression of 42 adhesion and signaling receptors, including 19 integrins, on a single-cell level. The NC cells had largely homogenous expression profiles, whereas the cancer cell lines showed a high degree of variation (**Extended data Fig.2a**). The integrins α6 and β4 cell surface expression levels were heterogenous, ranging from high to very low, in cancer cells compared to NC cells based on mass cytometry analysis (**Fig. 2c; Extended data Fig.2b**) and confocal immunofluorescence imaging (**Fig. 2d**). Staining of α6β4-associated hemidesmosome components BP180 (COLXVII) and keratin 14 reflected a similar heterogeneity and indicated a clear overall loss of hemidesmosomes and their associated intermediate filament cytoskeleton in the T3 cancer cells. Similar changes were also detected on bulk mRNA and protein levels of α6, β4, BP180 and keratin 14 (**Extended data Fig.2c-e**). Cell-surface expression of integrins α3 and β1 was also heterogeneous in cancer cells (**Fig. 2e**) and confocal immunofluorescence imaging demonstrated that this was linked to a striking difference in subcellular integrin localization rather than absolute changes in protein expression (**Fig. 2f**). Integrin α3 unexpectedly localized predominantly in cell-cell junctions in NC and T1 cells, whereas junctional localization was significantly decreased, and shifted to endosome-like intracellular structures in T3 cancer cells (**Fig. 2g**). The same was evident for the tetraspanin CD151, which interacts with α3β1 integrin with high affinity, localizing to focal contacts and hemidesmosomes^42,43^ (**Fig. 2h**). Furthermore, the cancer cells had an increased number of smaller vinculin-, active integrin β1 (12G10)- and integrin-linked kinase (ILK)-positive cell-matrix adhesions compared to NC cells (**Fig. 2i & j; Extended data Fig.2g & h**). Intriguingly, in addition to junctional localization, integrin α3 also localized in cryptic lamellipodia, which regulate epithelial cell migration^44^, in NC and T1 cells (**Fig.2g**). These marked changes in laminin-binding integrins imply that cell-cell and cell-matrix adhesion are altered in VFC.

**Fig. 2.**
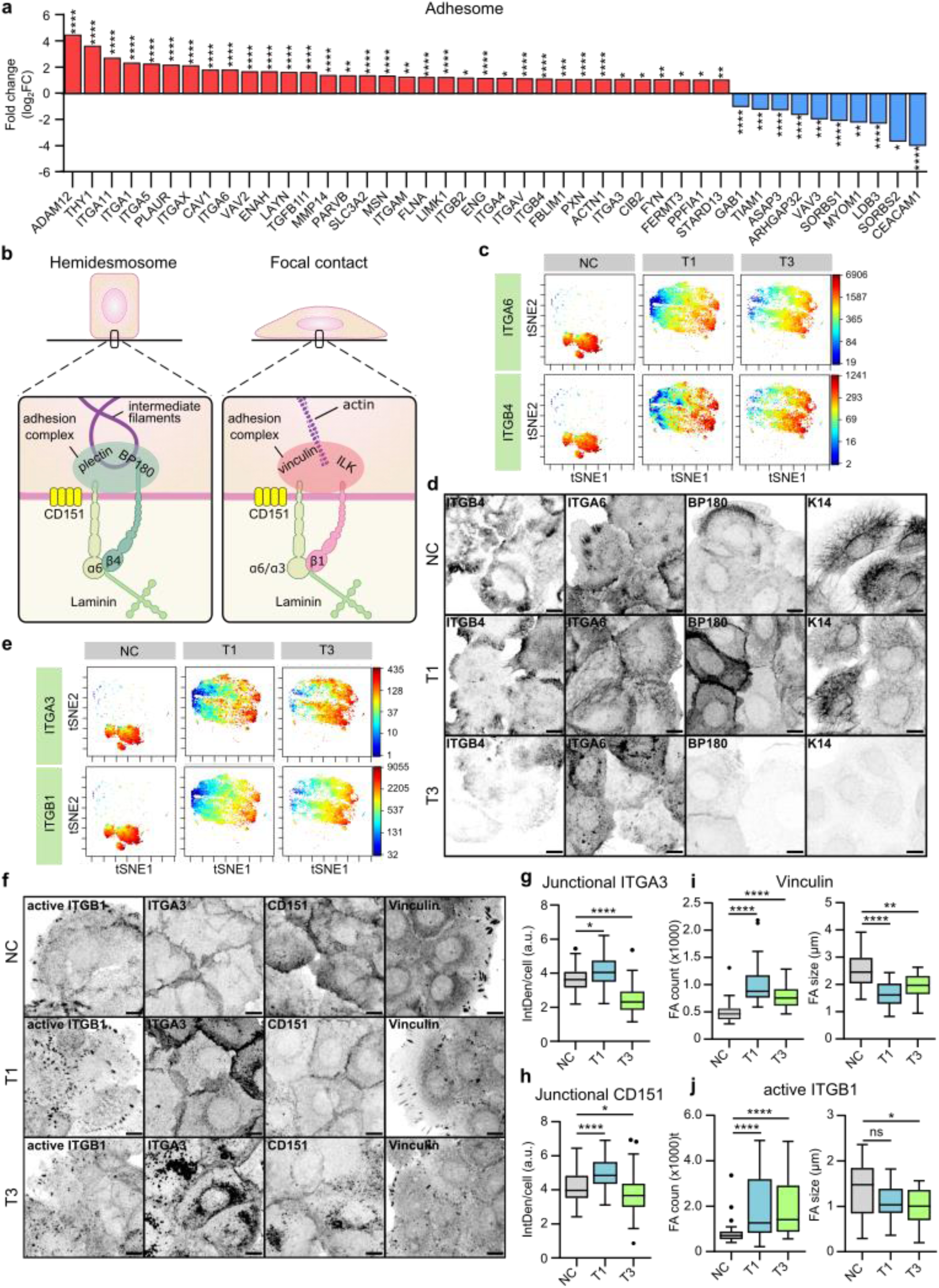
**Expression and subcellular localization of laminin-binding integrins is altered in vocal fold cancer a**, Differentially upregulated and downregulated (fold change, log2) adhesome^38,39^ genes in VFC (T1-T4, n=54) compared to normal (n=12) patient tissue (TCGA-data, FDR < 0.05). **b**, Schematic of laminin-binding integrins in hemidesmosomes (α6β4) and focal contacts (α6β1 and α3β1) connecting epithelial cells to the keratin cytoskeleton via BP180 and plectin or actin cytoskeleton via ILK and vinculin. **c**, t-distributed stochastic neighbor embedding (t-SNE) visualization of ITGA6 and ITGB4 single-cell surface expression (MassCytof) in NC cells and vocal fold T1 and T3 cancer cells. **d**, Representative ITGA6, ITGB4, BP180 and K14 confocal immunofluorescence images of NC cells and VFC T1 and T3 cells (n=3). Scale bar 10 μm. **e**, t-SNE visualization of ITGA3 and ITGB1 single-cell surface expression (MassCytof) in NC cells and vocal fold T1 and T3 cancer cells. **f**, Representative ITGA3, active ligand-engaged ITGB1 (12G10), CD151 and vinculin confocal immunofluorescence images of NC cells and vocal fold T1 and T3 cancer cells (n=3). Scale bar 10 μm. **g & h**, Quantification of junctional ITGA3 (**g**) and CD151 (**h**) in NC (ITGA3 n= 200, CD151 n=209) cells and vocal fold T1 (ITGA3 n=200, CD151 n=199) and T3 (ITGA3 n=199, CD151 n=205) cancer cells. **i & j**, Quantification of FA number (count) (left) and size (right) using vinculin (**i**) and active ITGB1 as markers in NC cells (vinculin n=29-30, ITGB1 n= n=28-30), and VFC T1 (vinculin n=30, ITGB1 n=30) and T3 (vinculin n=30, ITGB1 n=29-30) cells. Data are mean box plots or tukey mean-difference plots. n is the total number of cells/ average FA count/size per cell in field of view (FOV) pooled from three independent experiments. FDR was used to asses statistical significance of differentially expressed genes and Kruskal-Wallis test followed by post hoc Dunn’s multiple comparisons test was used to asses statistical significance of junctional and FA proteins.

### Stiffening of vocal fold tissue supports increased cell proliferation, migration and invasion

As we detected an increase in patient tissue stiffness and ECM expression in cancer, we set out to determine whether changes in stiffness influence VFC cell proliferation. We monitored cell proliferation for 4 days on collagen I and fibronectin- or Matrigel- (mainly composed of laminin and collagen IV) coated hydrogels of varying stiffnesses (0.5 kPa, 25 kPa and 50 kPa) and on plastic. T3 cell proliferation on collagen I and fibronectin-coated plates was significantly higher than those of T1 cells. Both T1 and T3 cells proliferated better on stiffer matrices (**Fig. 3a-b; Videos 1-6**) with more active β1-integrin in adhesions and better cell spreading on stiff **(Extended data Fig.3d**). Similar data were obtained on Matrigel-coated plastic and hydrogels (**Extended data Fig.3a-c; Extended data Fig.3e; Videos 7-12**). As single cells, T3 cells demonstrated increased speed, accumulated distance and directionality compared to T1 cells on collagen I and fibronectin-coated 50 kPa hydrogels (**Fig. 3d & e)**. Moreover, T3 collective cell migration (as a sheet in wound healing experiments) was significantly faster compared to T1 cells both on collagen I and fibronectin- (**Extended data Fig.3f-g**) and Matrigel-coated plastic plates (**Extended data Fig. 3h-i**). Accordingly, T3 cells invaded effectively through Matrigel transwell inserts (45h), whereas only a small number of T1 cells were able to invade (**Fig. 3f & g**). Taken together, these data indicate VFC proliferation and migration are positively regulated by increased ECM rigidity.

**Fig. 3.**
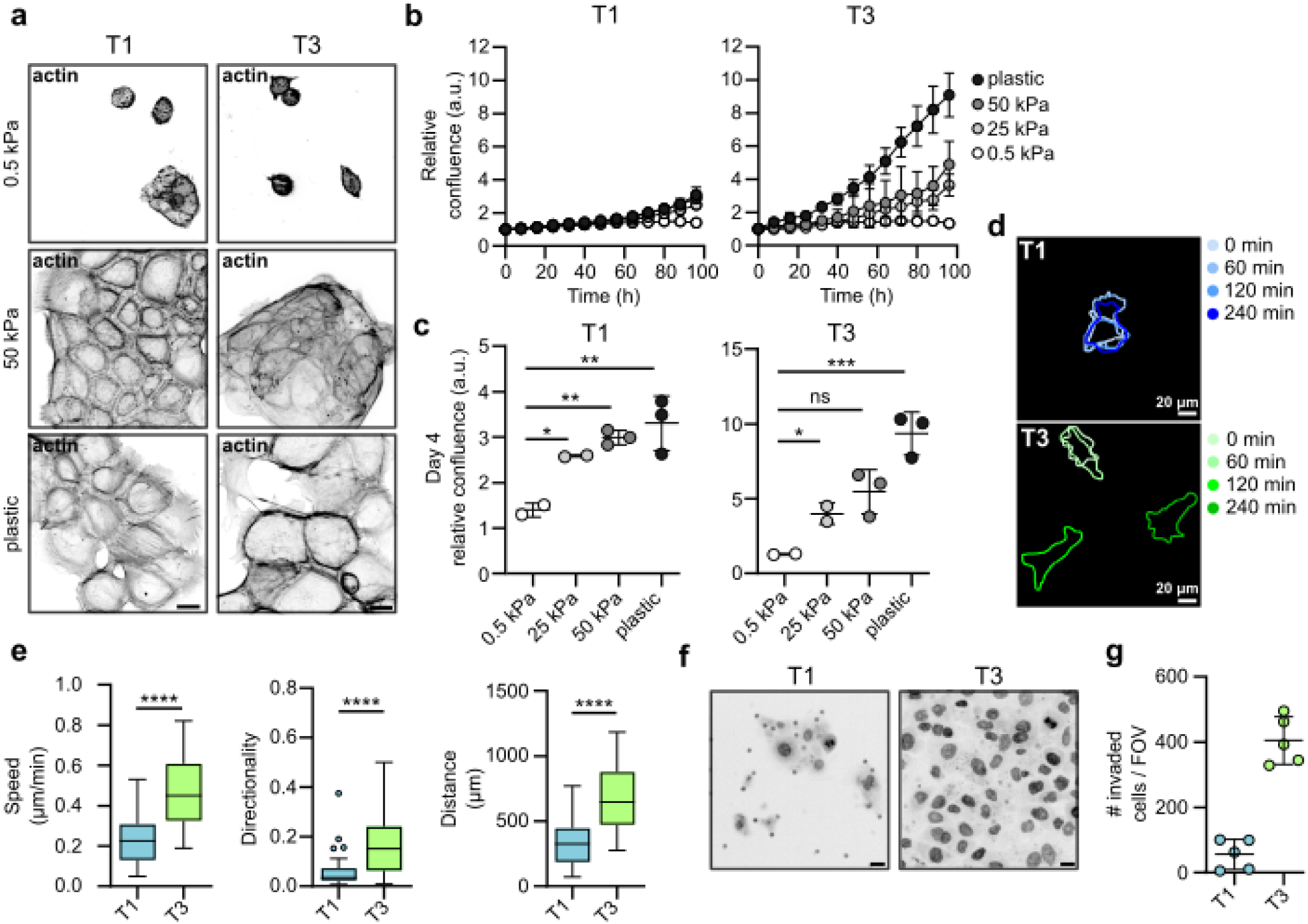
Stiffening of vocal fold tissue supports increased cell proliferation, migration and invasion a, Representative actin confocal immunofluorescence images of T1 and T3 VFC cells on 0.5 and 50 kPa hydrogels and plastic coated with collagen I and fibronectin (n=3). Scale bar 50 μm. **b & c**, Proliferation (**b**) of T1 and T3 VFC cells on hydrogels of varying stiffnesses (0.5 kPa, 25 kPa, 50 kPa) and plastic and confluence at end-point (**c**). **d & e**, Representative outlines (**d**) of T1 and T3 VFC single-cell migration on 50 kPa hydrogels at different timepoints (0 min, 60 min, 120 min and 240 min) and quantification (**e**) of speed (μm/min), distance (μm) and directionality (n=2). **f**, Representative nuclei (dapi) confocal immunofluorescence images (transwell pores visible in images as dots) and number of invaded T1 and T3 VFC cells per FOV in a Matrigel invasion assay (45h). Scale bar 20 μm. (n=2). Data are mean (± s.d.) or tukey box plots. Statistical significance was assessed using Kruskal-Wallis test followed by post hoc Dunn’s multiple comparisons test or Mann-Whitney test.

### Inhibition of laminin-binding integrins modulates monolayer dynamics and disrupts cell clustering in 3D-spheroids

α3β1-integrin localization to cell-cell junctions in normal squamous cells was reported more than two decades ago^45^. While the role of α3β1-integrin in mediating cell-matrix adhesion and controlling cell polarity in stratified epithelia is well-established in vitro and in vivo^46^, the functional role of this receptor in intercellular adhesion of epidermal squamous cells has been controversial and the molecular details remain elusive^47^. To explore the functional role of laminin-binding β1-integrins in VFC, we treated cells with integrin α3- (P1B5), α6- (P5G10) and β1 (mAb13)-blocking antibodies. Live-cell imaging of sparse cell clusters revealed retraction of junctional and cell-edge lamellipodia with a concomitant slowing of cell movement in all cell lines, most notably NC cells, after dual inhibition of integrins α3 and α6 (**Videos 13-15**). Blocking E-cadherin had the opposite effect; weakened cell-cell adhesions supported the scattering of cell colonies by reducing cell-cell coordination and increasing cell elongation and movement (**Videos 16-18**).

In a 3D-spheroid model, blocking the subunits of laminin-binding integrins; the common β1 subunit, α3 alone or in combination with integrin α6 all resulted in increased spheroid area primarily in NC and T1 cancer cells when compared to IgG control (**Fig. 4a & b**). The observed increase in size was due to reduced spheroid compaction and significantly more dissociated cells (**Extended data Fig.4a**). These data imply a functional role for integrins in the cell-cell junctions in NC and T1 cells (**Fig. 2f & g**). The T3 spheroids grew rapidly into large spheroids and integrin inhibition did not trigger marked spheroid dissociation, concordant with intracellularly localized integrins.

**Fig. 4.**
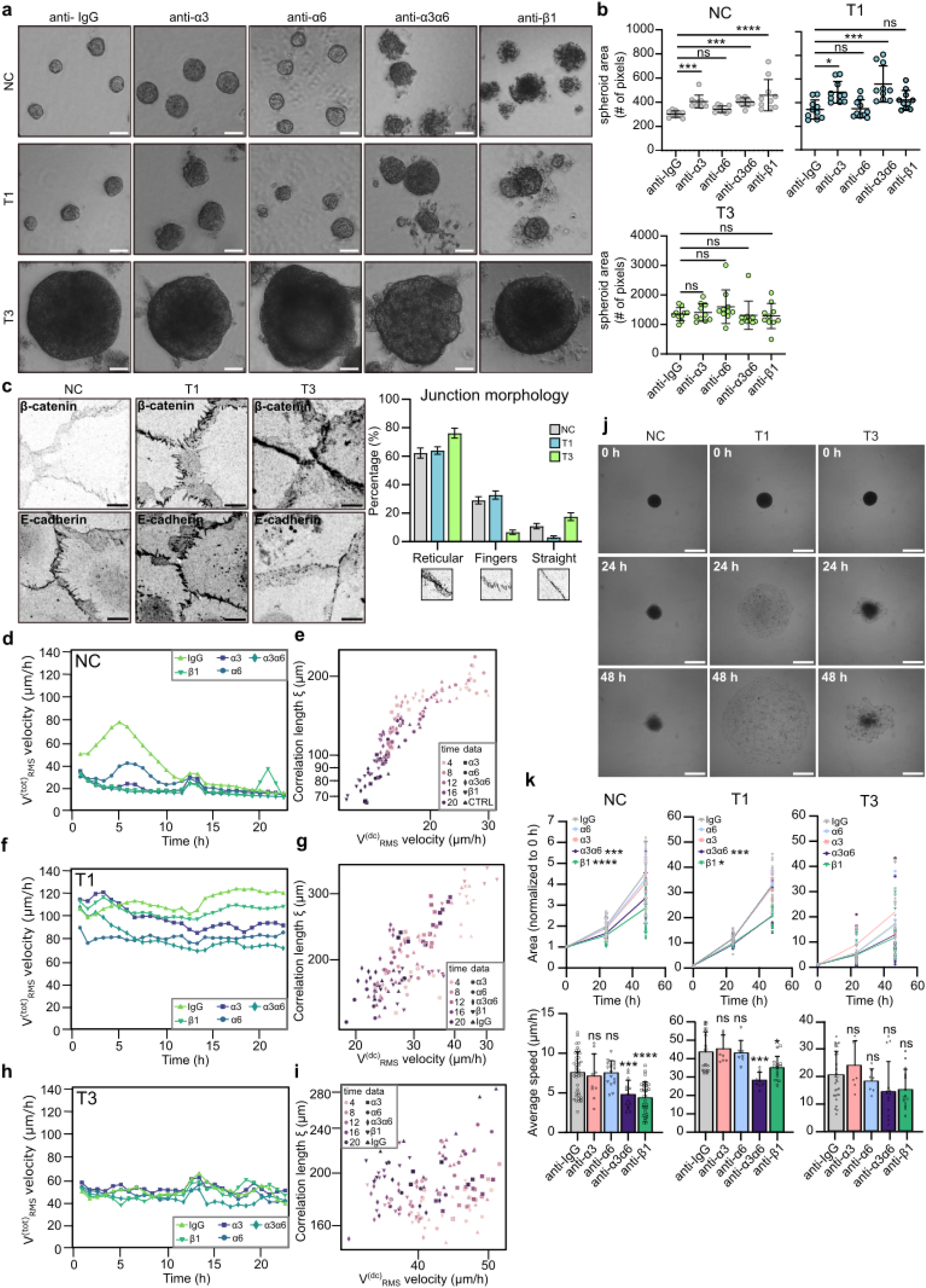
**Inhibition of laminin-binding integrins modulates monolayer dynamics and disrupts cell clustering in 3D-spheroids a & b**, Representative phase contrast images (**a**) and quantification (**b**) of spheroid size of NC cells and VFC T1 and T3 cells in 3D Matrigel cultures treated with IgG-control or integrin blocking antibodies (anti-α3, anti- α6, anti-α3α6 and anti-β1) for 11 days (n=3). Scale bar 50 μm. **c**, Representative β-catenin and E-cadherin confocal immunofluorescence images and quantification of junction morphology of NC and VFC T1 and T3 cells (n=3). Scale bar 10 μm. **d-i**, Quantification of total RMS velocity (d) and correlation length of NC (**d & e**) and VFC T1 (**f & g**) and T3 (**h & i**) cells treated with IgG-control or integrin blocking antibodies (anti-α3, anti-α6, anti-α3α6 and anti-β1) for 24h. **j & k**, Representative phase-contrast images (**j**) and normalized area and average speed (μm/h) (**k**) of NC and VFC T1 and T3 cells treated with IgG-control or integrin blocking antibodies (anti-α3, anti-α6, anti-α3α6 and anti-β1) undergoing wetting (n=3). Data are mean box plots (± s.d.). Statistical significance was assessed using Kruskal-Wallis test followed by post hoc Dunn’s multiple comparisons test.

These data prompted us to investigate VFC cell-cell junctions in more detail. T3 cells exhibited straight junctions (E-cadherin and β-catenin immunofluorescence staining), indicative of less tensile adhesions, whereas NC and T1 cells had protrusive finger-like junctions, indicative of more tensile adhesions (**Fig. 4c**). To quantitatively capture these differences, we divided junctions into three categories (straight, reticular and finger-like) based on morphology. Most notably, while reticular adhesions were a prominent feature in all cells, there was a near absence of finger-like-junctions and a larger proportion of straight junctions in T3 cells compared to NC cells and T1 VFC cells (**Fig. 4c**). Overall, these data indicate that cell-cell junctions are altered in VFC cell lines and that integrins contribute to junctional dynamics in NC and T1 VCF.

### VFC cells exhibit a previously unobserved solid-like flocking state ensuring long-range motility

Cell-cell and cell-matrix adhesions are critical determinants of the mechanics and dynamics of multicellular, normal and tumorigenic cell assemblies. At a critical cell density, motility of normal epithelia ceases and cells undergo a jamming phase transition (PT) which is considered a tumor- suppressive mechanism^48,49^, whereas PTs through unjamming and flocking motion, in turn, have been shown to promote collective modes of cancer invasion^50–53^. Thus, we next investigated monolayer dynamics of NC and VFC cells and the impact of integrin inhibition. PIV (Particle Image Velocimetry; see Materials and Methods for details) analysis revealed that untreated NC cells exhibited a progressive reduction in cell motility, quantified by the Root Mean Square velocity *v*^*tot*^_*RMS*_. (Fig. 4d). We also characterized the jamming transition by extracting the velocity correlation length *ξ* (expressing the size of a cluster of cells moving together), as well as the drift-corrected total RMS velocity *v*^*dc*^_*RMS*_ (*t*) (**Fig. 4e; Video 19**), used to isolate the disordered velocity component, minimizing the effects of drifts. NC monolayers show for all treatments the expected behavior i.e. initially large *ξ* and RMS velocities that simultaneously decrease over time across the jamming transition^54,55^. Inhibiting α3 (P1B5), α6 (P5G10) and β1 (mAb13) integrins significantly and robustly reduced the collective motion, resulting in an accelerated transition toward a jamming state, characterized by a progressive loss of degree of alignment in the cell velocity (**Extended data Fig.4b; Videos 20-23**).

Similar analyses were conducted on T1 and T3 cells. In both cases, the total RMS velocity (**Fig. 4f & h; Videos 24 & 29**) remained constant in time with values consistently larger than the final velocity for the NC cells. For T1 cells, inhibition of integrin b1 or integrins α3 and α6 together, were most efficient in reducing the total RMS velocity, suggesting a relevant role for these integrins in collective cell motility. In contrast, T3 cell motility was insensitive to integrin inhibition. Plotting the velocity correlation length *ξ* vs the drift-corrected total RMS velocity, *v*^*dc*^_*RMS*_ (*t*)(**Fig. 4g & I; Videos 25-28 & 30-33**) revealed a complete loss of correlation for T3 cells, and an intermediate behavior for T1 cells, suggesting that in both cases the tissues are far from a dynamically arrested, jammed state. Consistently, T1 VFC cells displayed cohesive and coordinated movement like bird flocking, with aligned cell velocities spanning the entire field of view (**Extended data Fig.4c**). Interestingly, these cells maintain long-range coordinated motion even when exposed to anti-integrin treatments. Similar flocking behavior was detected in the T3 cells, albeit to a lesser extent (**Extended data Fig.4d**). The absence of mutual cell rearrangements in VFC collective motility point to a mode of PT via a flocking solid transition, characterized by long-range coordinated motility in the absence of local cell rearrangements. Interestingly, flocking solid transition has been predicted by numerical simulation but has thus far not been observed experimentally in mammalian cells^56,57^. Collectively, our data suggest that VFC cells exploit a solid flocking-state to enhance long-distance collective motion, possibly contributing to invasion and metastasis in the cancer setting^58^. However, this remains to explored in future studies.

In keeping with this finding, we directly tested the ability of NC and VFC 3D spheroids to spread and diffuse onto ECM-coated substrate by undergoing a “wetting” transition^59–63^. This assay is thought to mimic the early step of local dissemination and depends on both the cohesive tensional state and viscoelastic properties of the cell aggregates and the cell-ECM interactions. Both T1 and T3 spheroids on FN-coated plates displayed a significant increase in wetting velocity compared to NC, but with a notable difference in morpho-dynamics. T1 spheroids rapidly wetted the surface, spreading with an elevated and uniform radial velocity consistent with the flocking solid mode of motion and elevated velocity correlation length *ξ* of the monolayer motility (**Fig.4j & k**). T3 spheroids, however, wetted the surface by extending irregular fronts, with protruding clusters and apparently contractile local regions, consistent with their high contractility and the reduced velocity correlation length *ξ* of the monolayer motility (**Fig.4j & k**). In NC spheroids, inhibition of α3 (P1B5), α6 (P5G10) and β1 (mAb13) integrins caused a notable reduction of wetting velocity. Conversely, only marginal effects on the wetting of both T1 and T3 spheroids were observed under these conditions (**Fig. 4j & k; Extended data Fig. 4e & f),** suggesting that VFC wetting was largely independent of cell-matrix adhesion receptors and likely dominated by the bulk mechanical properties of the 3D spheroids.

### Mechanical stimuli induce cytoskeletal and junctional alterations and cell extrusion in VFC

Prompted by the striking cell-intrinsic differences in the adhesive and mechanical properties observed between VFC and NC cells, we sought to determine if these alterations extended to the cellular response to mechanical stimuli. To recapitulate the mechanical forces in the vocal folds, we subjected the cells to two types of mechanical stimuli: stretching to mimic opening and closing of the vocal folds, and vibration, which occurs during phonation. Uniaxial cyclic stretching of cells (1Hz, 20% stretch) for 1 hour induced alignment (coherency) of the NC and T1 cancer cells perpendicularly to the stretch direction as exemplified by the visualization of actin filaments and phosphorylated myosin light chain (pMLC) (**Fig.5a & b, Extended data Fig.5a-b).** The poorly organized T3 cell monolayers did not show visible alignment, albeit their actin alignment (coherency) was significantly increased similarly as in NC and T1 cells (**Fig. 5b**). For the vibration, we chose a stimulus matching the frequency of human adult vocal fold during normal phonation^64^ (50-250 Hz, 1 min off/on). This induced actin stress fibers (**Fig.5c**) and caused marked remodeling of the monolayer. Furthermore, continued vibration for 6 hours induced a significant increase in extrusion of highly contractile, pMLC-positive cells in the T3 VFC, but not in the NC or T1 cells (**Fig. 5e-g; Extended data Fig.5c**). This suggests that vocal fold-like mobility in the T3 cell layer induces extrusion of cells akin to ejection of cells from crowded epithelia as a mechanism to ensure epithelial homeostasis and epithelium integrity^65^.

**Fig. 5.**
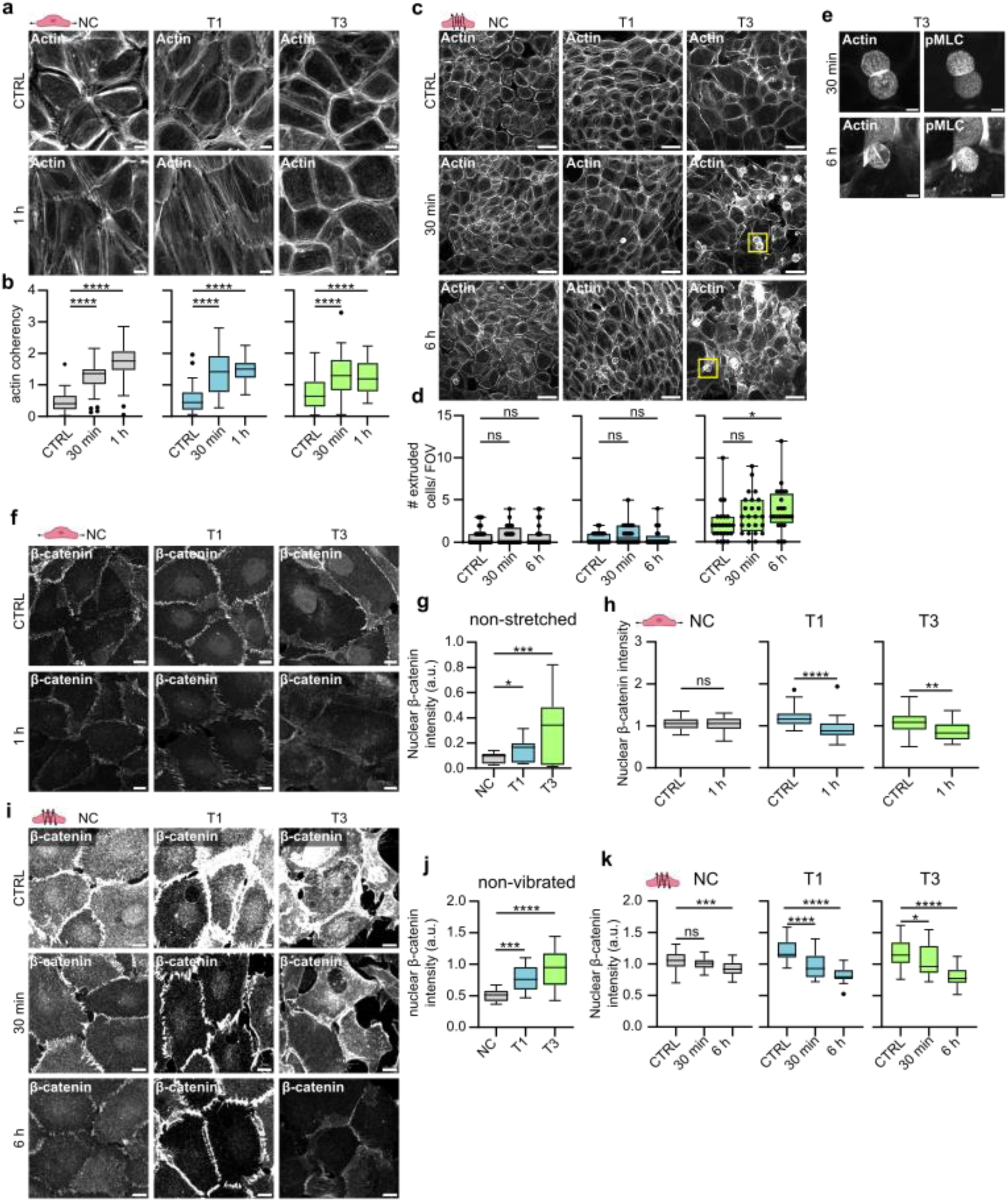
**Mechanical stimuli induce cytoskeletal and junctional alterations and cell extrusion in VFC a**, Representative actin confocal immunofluorescence images of NC cells and VFC T1 and T3 cells subjected to stretching (n=3). Scale bar 10 µm. **b**, Quantification of actin coherency (alignment) in stretched NC cells and vocal fold T1 and T3 cells (n=3). **c & d**, Representative actin confocal immunofluorescence images (**c**) and quantification of extruded (**d**) NC cells and VFC T1 and T3 cells subjected to vibration (n=3). Scale bar 50 µm. **e**, Representative actin and pMLC confocal immunofluorescence images of extruded T3 VFC cells subjected to vibration (n=3). Scale bar Scale bar 10 µm. **f-h**, Representative β-catenin confocal immunofluorescence images (**f**) and quantification of nuclear expression (integrated density per number of nuclei in FOV) of NC cells and VFC T1 and T3 cells in non-stretched conditions (**g**) and subjected to stretching (**h**) (n=3). Scale bar 20 µm. **i-k**, Representative β-catenin confocal immunofluorescence images (**i**) and quantification of nuclear expression (integrated density per number of nuclei in FOV) of NC cells and VFC T1 and T3 cells in non-vibrated conditions (**j**) and subjected to vibration (**k**) (n=3). Scale bar 20 µm.

Next, we investigated whether mechanical manipulation would cause changes in cell-cell junctions. Prior to stimulation, we noticed that β-catenin was significantly more nuclear in T1 and T3 cells compared to the NC cells (**Fig.5f-g**). This was particularly interesting, since nuclear β-catenin acts as a transcription factor activating signaling pathways that promote tumor formation^66,67^. Uniaxial cyclic stretching (1 hz, 20% stretch) for 1 hour caused alignment of β-catenin-positive junctions in NC and T1 cells (**Extended data Fig.5d**), and a significant reduction in nuclear and total β-catenin levels in the T1 and T3 cells (**Fig. 5h & i; Extended data Fig.5e**), which was also evident in vibrated cells (**Fig. 5j & k; Extended data Fig.5f**). Collectively, these data indicate that the cellular mechanoresponses under cyclic uniaxial stretch or vibration are different between NC and VFC cells, and mechanical stimulation of T3 cells, which represent the mechanically immobile stage of VFC in vivo, triggers cell extrusion and downregulation of oncogenic nuclear β-catenin.

### Phonomimetic mechanical stimuli decreases nuclear and total YAP levels

In addition to β-catenin, another key mechanosensitive oncoprotein in cancer is Yes-associated protein (YAP), which shuttles between the cytoplasm and nucleus, where it can activate downstream signaling pathways that maintain oncogenic signaling cascades^68^. Total YAP RNA (**Fig.6a**) and protein (**Fig.6b-c**) expression levels showed no significant changes in VFC cell lines compared to NC cells, but RNA expression of YAP downstream targets Cysteine-rich angiogenic inducer 61 (CYR61), Ankyrin Repeat Domain 1 (ANKRD1), AXL and macrophage colony stimulating factor (CSF1) were increased in VFC cells (**Fig.6d**), suggesting elevated pathway activity. Importantly, similarly to β-catenin, vibration decreased total and nuclear YAP levels in a time-dependent manner with prolonged vibration (6 hours) having a more significant effect than the acute 30 min stimulation. Concordant with these kinetics, the effect on the nuclear to cytoplasmic ratio, which is under acute mechanical control in many cell types, was less prominent and not significant in the T3 cells (**Fig. 6e-g and Extended data Fig.6a**). These data imply that cell vibration primarily regulates YAP levels rather that YAP mechanoresponsive shuttling to the nucleus.

**Fig. 6.**
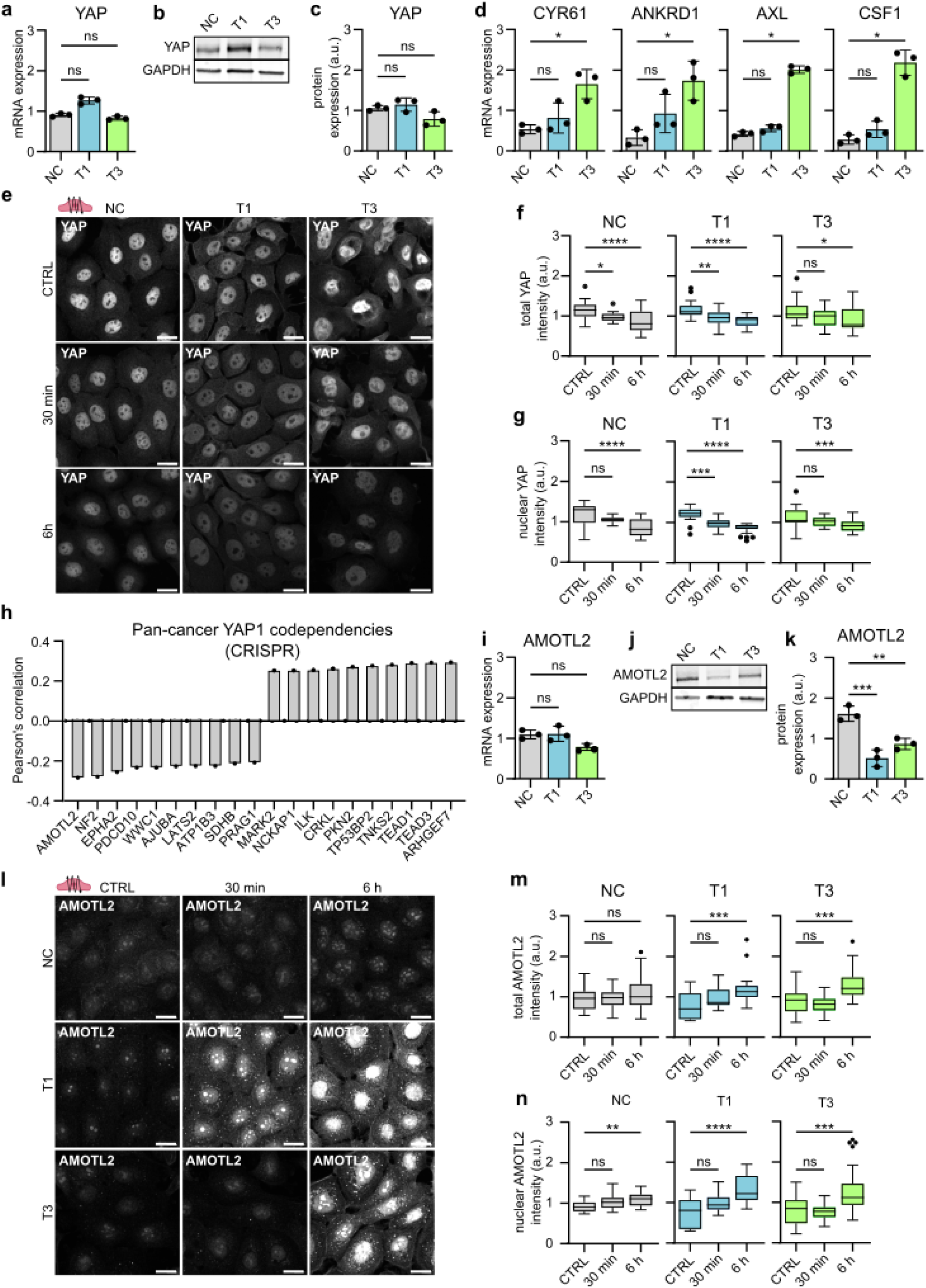
**Phonomimetic mechanical stimuli decreases nuclear and total YAP levels a**, Quantification of relative YAP mRNA expression (gene count) in NC cells and VFC T1 and T3 cells (n=3). **b & c**, Representative immunoblot (**b**) and quantification (**c**) of relative YAP protein expression in NC cells and VFC T1 and T3 cells (n=3). **d**, Quantification of relative RNA expression of YAP target genes CYR61, ANKRD1, AXL and CSF1 in NC cells and VFC T1 and T3 cells. **e**, Representative YAP confocal immunofluorescence images of NC cells and VFC T1 and T3 cells subjected to vibration (50-250 Hz, 1 min on/off) for 30 min or 6h compared to non-vibrated control (n=3). Scale bar 20 μm. **f**, Quantification of total (**f**) and nuclear (**g**) YAP intensity (integrated density) in NC cells and VFC T1 and T3 cells subjected to vibration (50-250 Hz, 1 min on/off) for 30 min or 6h compared to non-vibrated control (n=3). **h**, Quantification of Pan cancer YAP1 CRISPR codependency (DepMap) as Pearson’s correlation. **i**, Quantification of relative AMOTL2 mRNA expression (gene count) in NC cells and VFC T1 and T3 cells (n=3). **j & k**, Representative immunoblot (**j**) and quantification (**k**) of relative AMOTL2 protein expression in NC cells and VFC T1 and T3 cells (n=3). **l**, Representative AMOTL2 confocal immunofluorescence images of NC cells and VFC T1 and T3 cells subjected to vibration (50-250 Hz, 1 min on/off) for 30 min or 6h compared to non-vibrated control (n=3). Scale bar 20 μm. **m & n**, Quantification of total (**m**) and nuclear (**n**) AMOTL2 intensity (integrated density) in NC cells and VFC T1 and T3 cells subjected to vibration (50-250 Hz, 1 min on/off) for 30 min or 6h compared to non-vibrated control (n=3). Data are illustrated as tukey box plots or mean box plots ± s.d. (average of 8 FOV’s pooled from three independent experiments). Ordinary one-way Anova followed by post hoc Dunnett’s multiple comparisons test or Kruskal-Wallis test followed by post hoc Dunn’s multiple comparisons test was used to asses statistical significance.

To further understand the role of YAP in squamous cell carcinoma, we surveyed YAP1 cancer dependency maps on DepMap^69^. A pan cancer search identifying the top 20 co-dependencies in the CRISPR DepMap Public 23Q2+Score Chronos dataset found the strongest dependency hits (Pearson’s correlation, r) with Rho Guanine Nucleotide Exchange Factor 7 (ARHGEF7, r=0.29), TEA Domain Transcription Factor 3 (TEAD3, r=0.29), TEA Domain Transcription Factor 1 (TEAD1, r=0.29), Tankyrase 2 (TNKS2, r =0.28) and Angiomotin-like protein 2 (AMOTL2, r= -0.28) (**Fig.6h**). Moreover, integrin-linked kinase, which had increased FA localization in cancer cells, was one of the top 10 positive dependency hits (ILK, r=0.26) (**Extended data Fig.2g & h**).

Intrigued by these findings we sought to investigate the relationship between YAP and AMOTL2 in our cell model. AMOTL2 is a negative YAP regulator and has been shown to directly interact with YAP, retaining it within the cytoplasm^70–73^. AMOTL2 RNA levels were not significantly different between the cell lines (**Fig.6i**). However, AMOTL2 protein levels were significantly lower in VFC cells compared to NC cells (**Fig.6 j & k**). Vibration significantly increased AMOTL2 total and nuclear levels in VFC cells (**Fig.6l-n; Extended data Fig.6b**), coinciding with the decreased YAP levels (**Fig. 6f-g**). In summary, these results suggest that mechanical stimulation may decrease oncogenic nuclear YAP levels through an AMOTL2-dependent regulatory mechanism and the findings further support the notion of vocal fold mechanics contributing to tissue homeostasis, and having anti-oncogenic effects in VFC.

**Fig. 7.**
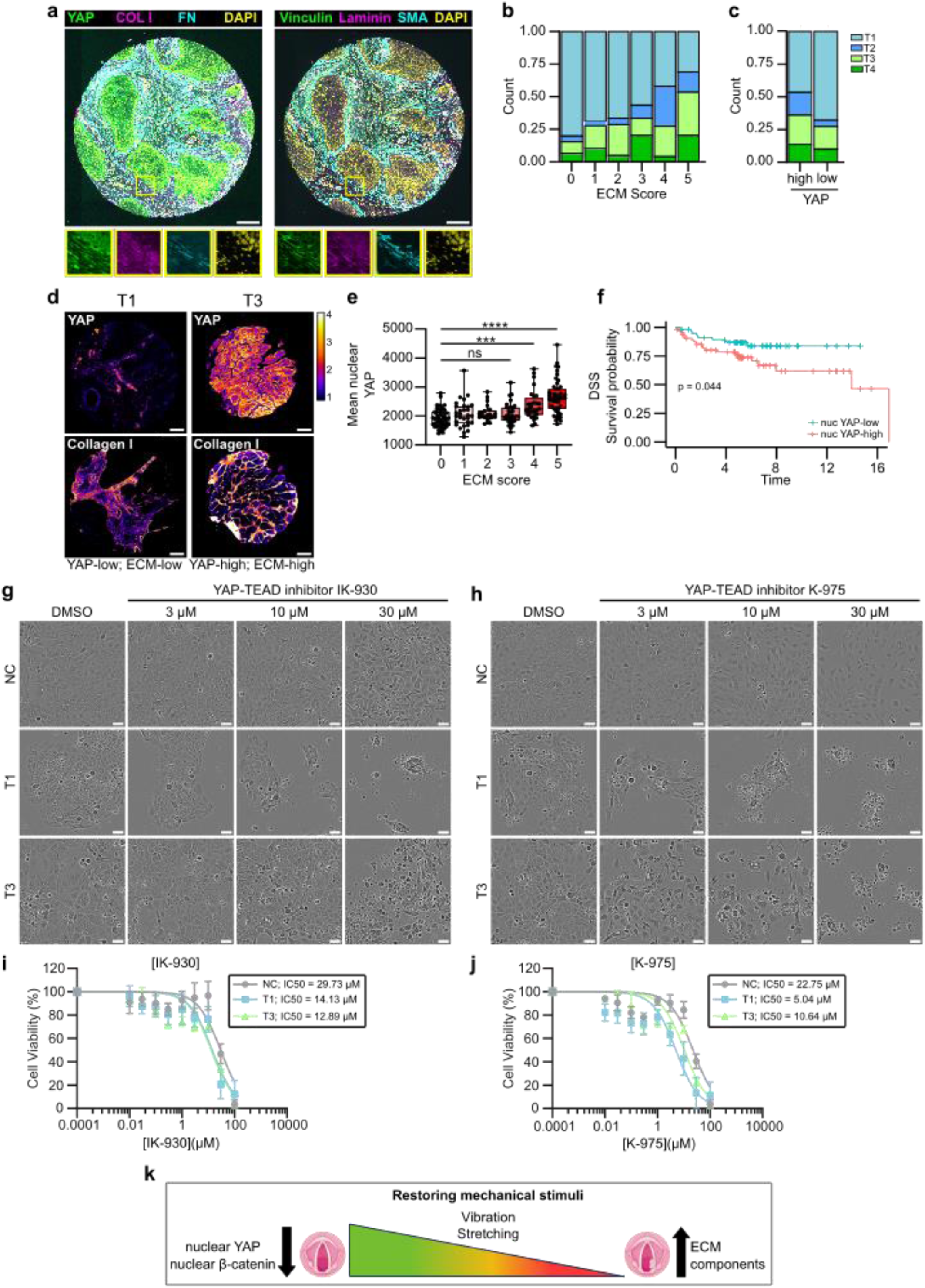
**High YAP levels correlate with high ECM expression and poor disease specific survival a**, Representative composite immunofluorescence images of TMA core stained for YAP/collagen-I/fibronectin/dapi or vinculin/laminin/SMA/dapi. Scale bar 100 μm. **b**, Quantification of correlation between ECM-score (median patient-level expression of stromal fibronectin, collagen-I, SMA, laminin and vinculin) and T-status in TMA multiplex histology. **c**, Quantification of correlation between YAP-score (median patient- level expression of epithelial YAP) and T-status in TMA multiplex histology. **d**, Representative YAP and collagen I staining of T1 and T3 cancer cells in YAP-low & ECM-low sample and YAP-high and ECM-high sample. Scale bar 100 μm. **e**, Quantification of correlation between ECM-score and mean nuclear YAP expression. **f**, Disease specific survival of YAP-high and YAP-low patients. **g & h**, Representative phase contrast images (**f**) and viability (**g**) of NC cells and vocal fold T1 and T3 cancer cells treated with YAP-TEAD inhibitor IK-930 for 48h. Scale bar 50 µm. **i & j**, Representative phase contrast images (**i**) and viability (**j**) of NC cells and vocal fold T1 and T3 cancer cells treated with YAP-TEAD inhibitor K-975 for 48h. Scale bar 50 µm. **k**, Graphical illustration of mechanical intervention of VFC cells as a therapeutic treatment option. Data are mean box plots (± s.d.). Statistical significance was assessed using Kruskal-Wallis test followed by post hoc Dunn’s multiple comparisons test or with Log-rank test for Kaplan-Meier analysis.

### High YAP levels correlate with high ECM expression and poor disease specific survival

To translate the in vitro findings into a more clinically relevant setting, we investigated the in vivo relevance of the identified mechanoregulators using multiplex histology and patient sample cohorts. We generated a custom laryngeal cancer tumor microarray (TMA) with cancer patient samples from T1 to T4 (n=218). We first noticed that there is a high correlation between all stromal ECM proteins (**Extended data Fig.7a**) and therefore implemented an ECM score, which considers median values for all the ECM and ECM-related proteins (FN, Col I, SMA, Laminin and Vinculin) in the tumor stroma across the patient cohort. Each patient was assigned an ECM score based on how many of the five ECM proteins were expressed at above-average levels, with scores ranging from 0 (all ECM proteins below average) to 5 (all ECM proteins above average). Scores of 0-2 were then classed as “ECM-low” while scores of 3-5 were classed as “ECM-high”. The analysis revealed a significant correlation between ECM score and T-status, with lower ECM scores being associated with lower T-status (**Fig.7a; Extended data Fig.7b-d**), but no significant correlation with patient survival (**Extended data Fig.7f**).

To determine whether YAP expression correlates with T-status (**Extended data Fig.7g&h**) and ECM score (**Extended data Fig.7h&j)** we calculated the median per-patient epithelial YAP-value in the dataset, and classified samples as either YAP-high or YAP-low based on this threshold. We found that YAP-high tumors tended to have higher staging, and patient-level nuclear YAP levels increased with higher ECM scores in tumors (**Fig.7b-d**). Moreover, we observed high YAP expression in T3-4 samples and identified nuclear YAP alone as being predictive of disease-specific survival (**Fig.7e**). Having established that patients with high ECM scores in the stroma have higher nuclear YAP in their tumor and a worse clinical outcome, we set out to explore whether inhibition of YAP-TEAD would affect cell viability. Treating cells with a YAP-TAZ-TEAD inhibitor, K-975, that covalently binds to a palmitate-binding pocket of TEAD and inhibits YAP function^74^, resulted in a significant and dose-dependent decrease in cell viability with the T3 VFC cells showing the highest sensitivity to the drug (**Fig.7f &g**). Another YAP-TAZ-TEAD inhibitor, IK-930, which is in phase I clinical trials for advanced solid tumors^75^ also showed increased sensitivity in VFC cells (**Fig.7h & i**). Taken together, these findings reveal clinical potential for YAP-TAZ-TEAD inhibition as a treatment option for VFC (**Fig.7j**).

### Outlook

Cells sense the biophysical features of their surrounding tissue, and the ensuing biomechanical signaling controls epithelial homeostasis, malignant progression, directed cell migration and drug sensitivity^13,76–78^. The vast majority of research in this area, however, draws from solid carcinomas arising from immobile tissue, such as the mammary gland, and the role of altered tissue mechanics in homeostasis and oncogenic properties of constantly moving epithelia remain poorly understood. Here we used cell culture models recapitulating key features of vocal fold epithelia including ECM rigidity, tissue stretching and vibration. We show that, concordant with the vocal fold epithelia becoming mechanically fixed and invasive with increasing T-status, VFC upregulates expression of multiple ECM components, is stiffer than normal vocal fold and proliferates in a stiffness dependent manner. Unlike kinetically arrested, densely packed (jammed) NC squamous epithelia, patient derived VFC cells are in a flocking, hyper-motile state, similar to the one previously established for invasive breast carcinomas^49,51^ in line with their high invasive capacity.

Cell cycle re-entry of arrested epithelia is regulated by nuclear translocation and transcriptional activity of YAP and β-catenin^79^. Malignant HNSCC tissues have higher YAP1 expression in comparison to benign patient samples, and YAP1 activation drives oral SCC tumorigenesis and correlates with poor patient survival^80–83^. However, YAP and β-catenin have not been explored in the molecularly distinct VFC^84^. We find that mechanical stretch and vibration, mimicking normal-like vocal fold mobility, downregulates nuclear β-catenin and nuclear YAP levels with a concomitant induction of the YAP inhibitor AMOTL2 in VFC cells derived from increasingly immobile tumors^85^. Moreover, high YAP correlates with a high ECM signature and poor clinical outcome in patient samples and VFC cells are increasingly sensitive to clinically tested^75^ YAP-TEAD small molecule inhibitors. Thus, normal tissue mechanics, mimicked in our cell culture systems by stretching and vibration, downregulate the activity of two relevant and synergistically acting oncogenic pathways^79^. These insights into the role of tissue mobility in maintaining homeostasis and suppression of malignancy may extend to other carcinomas arising from mobile epithelia and broaden our horizon on mechanical control of cancer progression.

## Material and methods

### The Cancer Genome Atlas (TCGA) data acquisition and analysis

The Cancer Genome Atlas (TCGA) HNSCC dataset was retrieved and filtered for patient ID’s with laryngeal cancer as tumor primary site. Pathology reports were then reviewed to asses tumor subsite and involvement of vocal folds. Raw files were downloaded from xena browser (https://xenabrowser.net/). Differentially expressed genes were assessed using Bioconductor R package ROTS (v.1.14.0), defining genes with FDR < 0.05 as differentially expressed^86^. Gene ontology was performed using ClusterProfiler (v. 4.8.3) in R^87^.

### Patient samples

Patient samples were obtained at the Department of Otorhinolaryngology-Head and Neck Surgery at Turku University Hospital under the Finnish Biobank Act with written informed consent from the sample donors (§279, 9/2001). Upon collection, the samples were given an arbitrary identifier and no patient identifiers, excluding patient age, and histopathological features of were available or recorded. Tissue samples were snap frozen with liquid nitrogen and stored at - 80°C until further processing.

### Atomic force microscopy

Atomic Force Microscopy (AFM) measurements of patient tissues were performed on freshly cut 16 µm snap frozen cryosections with JPK NanoWizard 4 (Bruker Nano) microscope mounted on an Eclipse Ti2 inverted fluorescent microscope (Nikon) and operated via JPK SPM Control Software v.6. Tissue sections were equilibrated in PBS with 1X protease inhibitors and measurements were performed within 30 minutes post thawing the sample. MLCT triangular silicon nitride cantilevers (Bruker) were used to access basement membrane stiffness. Forces of up to 3 nN were applied at 20 micron per second constant cantilever velocity. All analyses were performed with JPK Data Processing Software v.6 (Bruker Nano) by first removing the offset from the baseline of raw force curves, then identifying the contact point and subtracting cantilever bending before fitting the Hertz model with correct tip geometry to quantitate the Young’s Modulus.

### Cell lines and culture

HaCat (human immortalized squamous cells, ATCC), UT-SCC-11 (T1 human glottic laryngeal cancer, Turku University hospital), UT-SCC-103 (T3 human glottic laryngeal cancer, Turku University hospital) cells were cultured in DMEM (Dulbecco’s modified Eagle’s medium, Sigma-Aldrich) supplemented with 10% FBS (Sigma-Aldrich), 2 mM l-glutamine (Sigma-Aldrich) and 1% MEM nonessential amino acid solution (Sigma-Aldrich) at +37 °C, 5% CO2. UT-SCC-11 and UT-SCC-103 cell lines generated at Turku University Hospital have undergone scientific evaluation by Auria Biobank with a positive decision of release (AB22-7195) to be used in the study. All cell lines were regularly tested for mycoplasma with MycoAlertTM Mycoplasma Detection Kit (LT07-418, Lonza) and MycoAlertTM Assay Control Set (LT07-518, Lonza) to ensure mycoplasma-free culturing. Cells were washed with Phosphate-buffered saline (PBS) (Gibco™) and detached enzymatically with 0.25% trypsin-EDTA solution (L0932, Biowest).

### Proliferation assay

Plastic (Corning) or Softwell® Easy Coat (Matrigen, stiffness range: 0.5 kPa, 25 kPa and 50 kPa) 24-well plates were coated with 10 μg/ml collagen I (C8919, Sigma) and 10 μg/ml fibronectin (341631, Sigma) diluted in PBS or 10 μg/ml growth factor reduced Matrigel (354230, Corning®) diluted in PBS, at +37 °C for 1 h. Coated plates were washed three times with PBS prior to seeding 10 000 cells in culture medium. Time-lapse live-imaging was performed using Incucyte S3^®^ or ZOOM Live-Cell Analysis System for 96h with 2h imaging intervals (10x objective). Medium was changed every second day.

### Migration assay

50 kPa Softwell® Easy Coat (Matrigen) 24-well plates were coated with 10 μg/ml collagen I (C8919, Sigma) and 10 μg/ml fibronectin (341631, Sigma) diluted in PBS, at +37 °C for 1 h. Coated plates were washed three times with PBS prior to seeding 1000 cells in culture medium. Time-lapse live-imaging was performed using Nikon Eclipse Ti2-E (10x/ 0.3 objective) for 24h with 10 min imaging intervals. Single-cell tracking was performed using TrackMate plugin in FIJI (National Institutes of Health; NIH).

### Invasion assay

200 000 cells were seeded in serum free medium on Matrigel transwell inserts (354480, Corning) and placed in culture medium. After 45h of invasion, uninvaded cells in the inner well were wiped off with cotton buds and invaded cells were fixed with 4 % PFA diluted in PBS for 10 min at RT. Inserts were washed 3 times with PBS and stained overnight with Dapi. Invaded cells were assessed by confocal imaging (3i Marianas CSU-W1; 20×/0.8 objective) and quantifying the number of invaded cells per field of view (FIJI).

### Viability assay

5000 cells were per 96-plate well in culture medium. DMSO (D265, Sigma) or YAP-TAZ-TEAD inhibitors K-975 (HY-138565, MedChemExpress) or IK-930 (HY-153585, MedChemExpress) were added at 10 nM, 30 nM, 100 nM, 300 nM, 1µM, 3µM 10 µM, 30 µM and 100 µM concentrations the following day. Relative cell viability was measured as absorbance at 450 nm after a 2-hour incubation with a cell counting kit at +37°C as per the manufacturer’s instructions (Cell Counting Kit 8, ab228554) 48 h after addition of inhibitor treatment.

### Western blotting

Cells were kept on ice and washed with cold PBS and lysed with heated (+90°C) TX- lysis buffer (50 mM Tris-HCl, pH 7.5, 150 mM NaCl, 0.5% Triton-X, 0.5% glycerol, 1% SDS, Complete protease inhibitor [SigmaAldrich], and phos-stop tablet [Sigma-Aldrich]). Lysed cells were scraped into an Eppendorf tube and boiled for 5 min at +90 °C followed by 10 min sonication and 10 min centrifugation at 13000 rpm at +4°C in a microcentrifuge. Protein concentrations were determined from the supernatant with DC Protein assay (Bio-Rad) as per the manufacturer’s instructions. Samples were boiled at +90 °C for 5 min prior to protein separation using precast SDS-PAGE gradient gels (4–20% Mini-PROTEAN TGX, Bio-Rad) and transferred onto nitrocellulose membranes with the semi-dry Trans-Blot Turbo Transfer System (Bio-Rad). Membranes were blocked with AdvanceBlock-Fluor blocking solution (AH Diagnostics) diluted 1:1 in PBS for 1h at room temperature (RT) and incubated over night at +4°C with primary antibodies diluted in AdvanBlock-Fluor blocking solution. Membranes were washed for 5 min three times with TBST (Tris-buffered saline and 0.1% Tween 20) and incubated 1:2500 with fluorophore-conjugated Azure secondary antibodies (AH Diagnostics) in blocking solution for 1 h at RT. Membranes were washed three times with TBST for 5 min at RT. Membranes were scanned using an infrared imaging system (Azure Sapphire RGBNIR Biomolecular Imager) and band intensities were analyzed using Image Studio Lite (Licor) by normalizing the signal to GAPDH or HSP70, which were used as a loading controls.

### Particle-image velocity analysis (PIV)

A custom PIV algorithm was developed in Python to measure cell velocities within monolayers and derive different indicators of cellular motility. Velocity fields were first extracted by processing sequences of images. In short, each image is divided in square regions of interest (ROI), for each ROI located at position 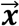 the local cell displacement 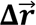 is quantified by cross correlating the intensity of two ROI-images separated by Δ*t*, which allows estimating the local velocity as 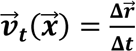, where the index *t* corresponds to the time of the frame pair used to compute the velocity field. We used ROIs of size 80 × 80 px^2^, which are slightly larger than the typically observed cell size of ∼ 50 px, with a spatial overlap factor between different ROIs of 50%. To improve statistics, we also performed a temporal average of the so-obtained velocity fields over chunks of length 20 frames (200 minutes), again with a temporal overlap of 50%. The previous parameters were carefully optimized to find the best tradeoff between increasing the spatiotemporal resolution and averaging a sufficient number of data samples to obtain smoother velocity maps, which will be indicated in the following with 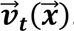. We then followed Garcia et al.^55^ to compute the total root mean square (RMS) velocitiy 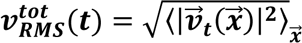 and the drift corrected RMS velocity 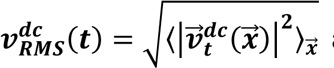 as spatial averages of the velocity fields, where we have introduced the drift collected velocity 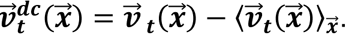. In cell lines with no strong collective motion, 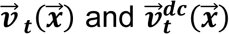 are similar, but in the presence of collective motion these two quantities can differ substantially. As suggested by Garcia et al.^55^, we used the drift-corrected velocity to calculate the radial velocity-velocity correlation function, obtained as

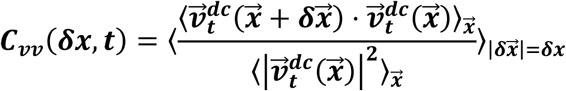

and we fitted this function to a model exponential 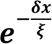 to extract the spatial correlation length ξ of the velocity field, quantifying the size of regions with similar velocities once the average monolayer velocity has been removed. Finally, to better visualize spatial correlations in the velocity field, we followed Malinverno et al.^56^ and calculated the alignment index 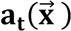 as the cosine of the angle between the average velocity vector of a single velocity field with every other velocity vector.

### Cell stretching assay

Stretch chambers (STB-CH-4W, STREX Cell Stretching Systems) were autoclaved and coated with 10μg/ml collagen I (C8919, Sigma) and 10μg/ml fibronectin (341631, Sigma) diluted in PBS at +37 °C for 2 h. Coated chambers were washed three times with PBS prior to seeding 200 000 cells per well in culture medium. Cells were stretched the following day with STREX cell stretching system (model # STB-140-10) with 20 % stretch (6.40mm), 1 Hz frequency for varying periods (5 min, 30 min 1 h).

### Cell vibration assay

Flexible-bottomed silicone elastomer plates (BF-3001U, BioFlex®) were coated with 10μg/ml collagen I (C8919, Sigma) and 10μg/ml fibronectin (341631, Sigma) diluted in PBS for 2h at +37 °C. Coated chambers were washed three times with PBS prior to seeding 500 000-900 000 cells in culture medium. On the following day, stimulation sound files were played for varying periods (5 min, 30 min 1h, 6h) 1 min off /1 min on at a frequency range of 50-250 Hz with a phonomimetic bioreactor^88^ connected to a Crown XLS 1502 amplifier.

### 3D spheroid assay

Spheroid formation in a 3D environment was assessed by embedding cells between two layers of Matrigel (Corning, 354230). Firstly, the bottom of an angiogenesis 96-well µ- plate (89646, Ibidi GmbH) was coated with 10 µl of 50% Matrigel diluted in culture medium and centrifuged at +4°C, 200 g for 20 min followed by 1-hour incubation at +37°C. Next, wells were filled with 20 µl of cell suspension in 25% Matrigel diluted in culture medium (500 cells/well), centrifuged for 10 min at 100 g and incubated at +37°C for 4h. Wells were filled with culture medium supplemented with 10 µg/ml function blocking antibodies or IgG control; mouse anti-IgG (31903, Invitrogen), mouse anti-human α3 integrin (P1B5, In-house hybridoma), mouse anti-human α6 integrin (P5G10, In-house hybridoma) and rat anti-human β1 integrin (mAb13, In-house hybridoma). Spheroid formation was imaged for 10 days with IncuCyte S3 Live-Cell Analysis system (10x objective). Culture medium was changed every 2-3 days. Analysis was performed using OrganoSeg software^89^ and ImageJ.

### Wetting assay

Cells were seeded in a low attachment round bottom 96-well plate to allow the formation of spheroids. The following day, spheroids were transferred to a multiwell plate previously coated with 10ug/ml Fibronectin (diluted in PBS, incubated overnight at +4°C, and washed twice with PBS). Spheroids were monitored as they wet the substrate by time lapse imaging for 48h using iXplore live Microscope (Olympus Evident) (4x Objective, 10 min timeframe). Analysis of spreading area over time was performed using ImageJ. The data were normalized to the area of the spheroid at time 0. To evaluate the impact of Integrin perturbations, spheroids were treated with the blocking antibodies described above before starting the wetting experiment.

### Immunostaining

Coated (as previously mentioned) µ-slide 8-well chambered coverslips (Ibidi), standard culture plates (Corning) or Softwell® Easy Coat (Matrigen) were fixed at indicated endpoint with 4% PFA in culture medium for 10 min at RT. Cells were washed with PBS three times for 5 min. Permeabilization and blocking was performed using 0.3% Triton-X-100 in 10% normal horse serum diluted in PBS for 20 min at RT. Cells were stained with primary antibodies diluted in 10% normal horse serum overnight at 4°C. Cells were washed three times for 5 min with PBS and incubated with secondary antibodies diluted in PBS for 1h at RT, followed by three 5-min washes with PBS. Samples were either imaged right away or stored at 4°C covered from light until imaging.

### Imaging

Confocal imaging was performed with a 3i spinning disk confocal (Marianas spinning disk imaging system with a Yokogawa CSU-W1 scanning unit on an inverted Carl Zeiss Axio Observer Z1 microscope, Intelligent Imaging Inno-vations, Inc., Denver, USA) with 10x Zeiss Plan-Apochromat objective (without immersion, 2mm working distance,0.45 numerical aperture), 40x Zeiss LD C-Apochromat objective (water immersion, 0.62mm working distance, 1.1 numerical aperture) and 63x Zeiss Plan-Apochromat objective (oil immersion, 0.19 mm working distance, 1.4 numerical aperture). Widefield imaging was performed with Nikon Eclipse Ti2-E (Hamamatsu sCMOS Orca Flash4.0, Lumencor Spectra X LED excitation). Live imaging was performed with Incucyte S3 or ZOOM Live-Cell Analysis System.

### Mass Cytometry

Cells were grown on a 10 cm plate to 90% confluence, washed once with PBS and detached with cell dissociation buffer (#13150-016, Gibco). Detached cells were dispensed into 15 ml falcon tubes, centrifuged at 300 x g for 5 min followed by removal of supernatant and mixing the pellet by pipetting. Cells were resuspended in 1 ml of serum free medium. 1 ml of 1µM cisplatin in serum free medium was added to cells for 5 min, mixed well by pipetting and incubated for 5 min at room temperature. The mixture was quenched with Cell Staining Buffer (Maxpar®), 5x vol of the stained cells. Cells were centrifuged at 300xg for 5 min, the supernatant aspirated and cells resuspended by pipetting. Cells were washed with 4 ml of Cell Staining Buffer (Maxpar®). Cells were counted and 3 million cells aliquoted into 5 ml polypropylene tube followed by centrifugation at 300xg for 5 min. The supernatant was aspirated and cells gently mixed by pipetting. Cells were resuspended in 50 ul of Cell Staining Buffer (Maxpar®). Cells were then stained with the antibody panel, starting with Fc-blocking. Fc Receptor Blocking Solution was added 1:100 to each tube and incubated 10 min at room temperature. 50 ul of the prepared antibody cocktail was added to each tube and gently mixed by pipetting and incubated at room temperature for 15 min. Samples were gently vortexed and incubated for an additional 15 min at room temperature. After a total of 30 min incubation, samples were washed by adding 2 ml Cell Staining Buffer (Maxpar®) to each tube, centrifuged at 300xg for 5 min and the supernatant was removed. Sample wash was repeated three times and cells were resuspended in residual volume by gently vortexing after final wash and aspiration. Cells were fixed with 1 ml of 1.6% FA diluted in PBS and gently vortexed before 10 min incubation at room temperature. Samples were centrifuged at 800x g for 5 min and the supernatant was removed. Samples were gently vortexed to resuspend in residual volume. After cell staining, 1 ml of cell intercalation solution was prepared for each sample by diluting Cell-ID Intercalator-103Rh 1:1000 into Fix and Perm Buffer (Maxpar®) and mixed by vortexing. 1 ml of intercalation solution was added to each tube and gently vortexed. Samples were incubated 1h at room temperature or left overnight at + 4°C (up to 48h). Before acquisition with Helios (WB Injector) cells were at 800 x g for 5 min and washed by adding 2 ml of Cell Staining Buffer (Maxpar®), followed by another round of centrifugation. The supernatant was removed and samples gently vortexed to resuspend cells in residual volume. Cells were washed by adding 2 ml of CAS to each tube and gently vortexed before counting and transferring 1 million cells into a new tube. Tubes were centrifuged at 800 x g for 5 min, followed by careful aspiration of supernatant. Cells were gently vortexed to resuspend in residual volume and finally 1 million cells were resuspended in 900 ul CAS. Cells were filtered into cell strainer cap tubes. Sufficient volume of 0.1X EQ beads to re-suspend all samples in the experiment were prepared by diluting 1-part beads to 9-parts CAS. Cells were left pelleted until ready to run on Helios. Immediately prior to data acquisition, cell concentration was adjusted to 1.0 x 106 cells/ml diluted EQ bead solution. Cells were filtered into cap tubes. Samples were run and data acquired with Helios CyTOF. Mass cytometry antibodies were either purchased from Fluidigm or self-conjugated.

### RNA-sequencing

RNA was isolated from three biological replicates of cells seeded on coated BioFlex® plates. Cells were washed with cold PBS followed by RNA extraction using NucleoSpin RNA -kit (#740955.250, Macherey-Nagel) as per the manufacturer’s instructions. Total RNA concentration was measured with Nanodrop and samples were normalized by diluting with RNAse free water. Sample quality was verified using Agilent Bioanalyzer 2100, and final concentrations were measured using Qubit®/Quant-IT® Fluorometric Quantitation (Life Technologies). Illumina stranded total RNA prep library was prepared using 100 ng of RNA) as per the manufacturer’s instructions (Illumina Stranded mRNA Preparation and Ligation kit, (Illumina) and sequenced with Novaseq 6000 (S4 instrument, v1.5 (Illumina), 2x50 bp, SP flow cell, 2 lanes (650-800 M reads). Library quality was verified using Advanced Analytical Fragment Analyzer. The sequencing data read quality was ensured using the FastQ (v.0.11.14) and MultiQC (v.1.5) tools^90^. Differentially expressed genes were assessed using Bioconductor R package ROTS (v.1.14.0) defining genes with FDR < 0.05 as differentially expressed.

### Tissue microarray (TMA)

TMA blocks with duplicate core biopsies were made from formalin-fixed, paraffin-embedded tissue samples using a TMA Grand Master (3DHISTECH, Budapest, Hungary) at Helsinki University hospital. A total of 218 patients with known TNM staging and survival end- points were included in the study.

### Primary antibodies

**Table 1.** list antibodies used for Mass Cytometry.

| Metal tag | Target protein | Conjugation |
| --- | --- | --- |
| 106CD | a11 integrin | Self-conjugated |
| 110CD | HER3 | Self-conjugated |
| 111CD | a3 integrin (CD49c) | Self-conjugated |
| 112CD | EGFR | Self-conjugated |
| 113CD | CD10 | Self-conjugated |
| 114CD | av integrin (CD51) | Self-conjugated |
| 116CD | HER4 | Self-conjugated |
| 89Y | allb integrin (CD41) | 3089004B |
| 141PR | EpCAM (CD326) | 3141006B |
| 142ND | PETA-3 (CD151) | 3142011B |
| 143ND | N-Cadherin (CD325) | 3143016B |
| 144ND | Syndecan-4 | Self-conjugated |
| 145ND | Syndecan-1 (CD138) | 3145003B |
| 146ND | b3 integrin (CD61) | 3146011B |
| 147SM | ALCAM (CD166) | Self-conjugated |
| 148ND | HER2 (ErbB2/EGFR2) | 3148011A |
| 149SM | CD34 | 3149013B |
| 150ND | avb3 integrin (CD51/61) | 3150026B |
| 151EU | ICAM-2 (CD102) | 3151015B |
| 152SM | avb5 integrin | Self-conjugated |
| 153EU | b6 integrin | Self-conjugated |
| 154SM | Notch1 | Self-conjugated |
| 155GD | a8 integrin | Self-conjugated |
| 156GD | b1 integrin (CD29) | 3156007B |
| 158GD | E-Cadherin (CD324) | 3158018B |
| 159TB | LAT1 (CD98) | 3159022B |
| 160GD | a5 integrin (CD49e) | 3160015B |
| 161DY | a2 integrin (CD49b) | 3161012B |
| 162DY | b7 integrin | 3162026B |
| 163DY | a1 integrin (CD49a) | 3163015B |
| 164DY | a6 integrin (CD49F) | 3164006B |
| 165HO | Notch2 | 3165026B |
| 166ER | CD44 | 3166001B |
| 167ER | Notch3 | Self-conjugated |
| 168ER | a9b1 integrin | 3168013B |
| 169TM | CD24 | 3169004B |
| 170ER | ICAM-1 (CD54) | 3170014B |
| 171YB | CD9 | 3171009B |
| 172YB | Neuropilin-1 (CD304) | Self-conjugated |
| 173YB | b4 integrin (CD104) | 3173008B |
| 174YB | a4 integrin (CD49d) | 3174018B |
| 175LU | b8 integrin | Self-conjugated |
| 176YB | NCAM (CD56) | 3176001B |
| 209BI | CD47 | 3209004B |

**Table 2:**
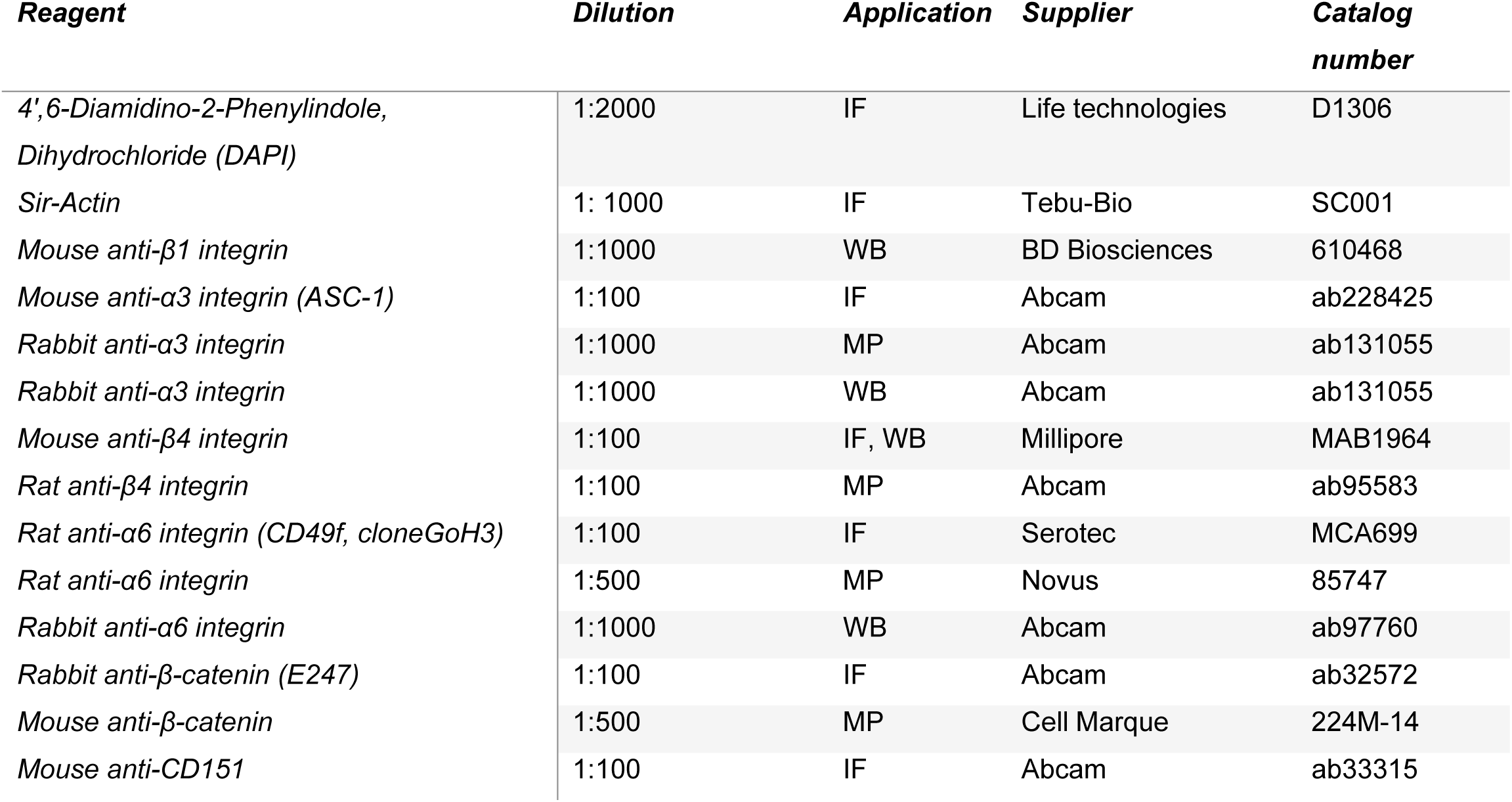
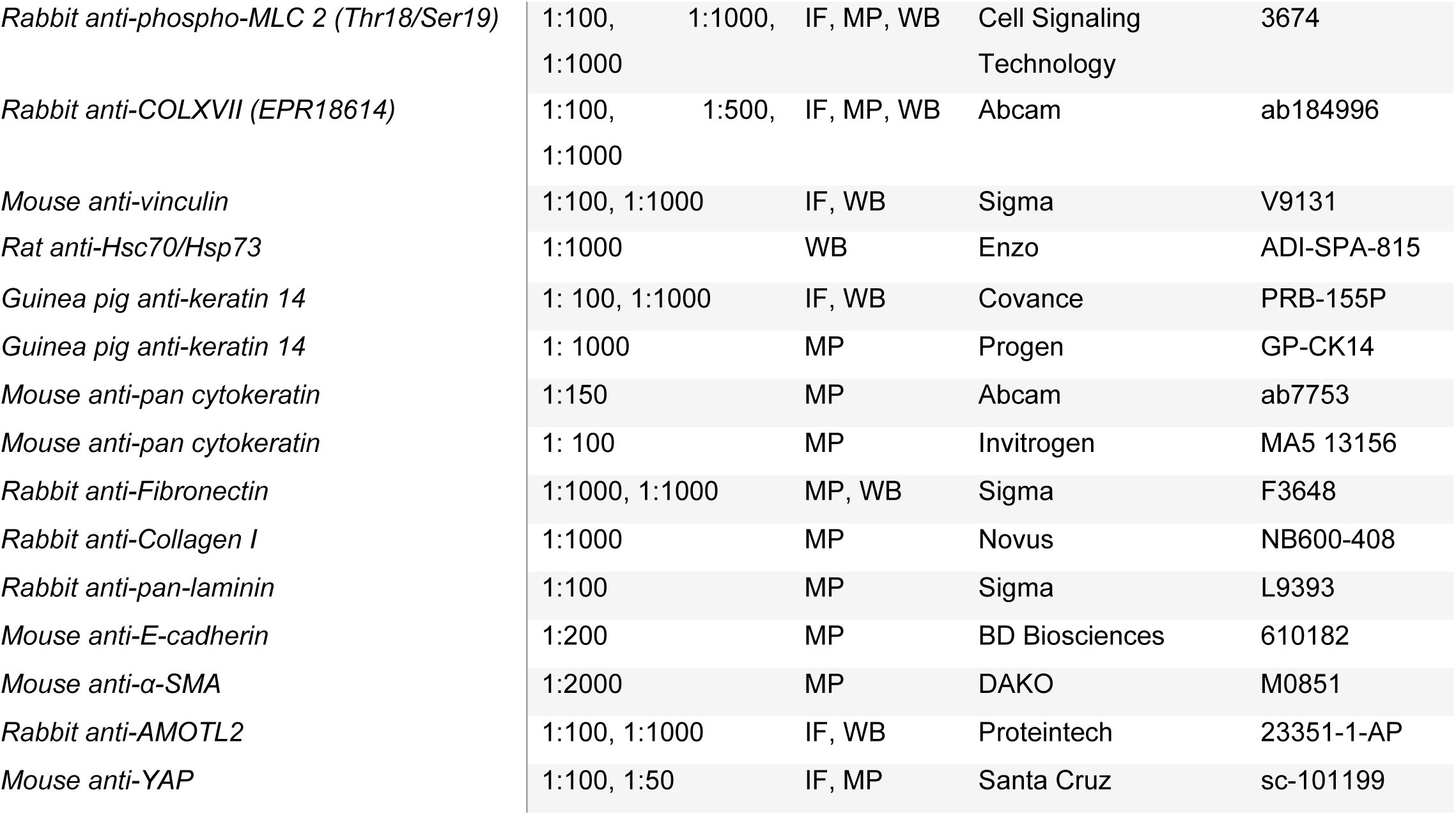
Details of primary antibodies used in the study. IF= immunofluorescence, MP= multiplex fluorescence immunohistochemistry, WB= western blot.

### Multiplexed fluorescent immunohistochemical staining and imaging

Multiplexed fluorescent immunohistochemical staining and imaging was performed in three cycles as previously described^91^ for two sets of seven to eight antibodies and the nuclear marker DAPI (Table 2), stained on two serial TMA sections. After the first-round staining and whole-slide imaging of the TMAs, the fluorescence signal was bleached, and the antibodies from the first-round staining were denatured, after which the second-round staining was performed. The process was repeated for the third round of staining. Imaging was performed using a Zeiss Axio Scan.Z1 slide scanner, with each round of staining recorded as an independent .CZI image file containing up to five fluorescent channels.

### Image analysis of multiplexed TMA datasets

Images of individual TMA cores were extracted from the whole-slide images using the TMA dearrayer functionality in QuPath^92^. Images from the three staining rounds were registered using an affine image registration method operating through the pyStackReg Python dependency^93^, aligning the DAPI channels of the three staining rounds. Autofluorescent signal from red blood cells and other histology artefacts (e.g. wrinkled or folded tissue section areas) were removed using a pixel classifier in Ilastik^94^. Nuclei were segmented from the DAPI channel using a trained StarDist model^95^. The nuclear regions of interest (ROIs) were expanded by 6 pixels to generate extra-nuclear ROIs. Pan-epithelial staining was used to threshold cells into epithelial and stromal compartments. A custom python script was then used to calculate fluorescence intensity in all channels for the relevant nuclear or extra-nuclear ROI in the relevant tissue compartments. Finally, patient-level average expression values were calculated for all cells and all TMA cores originating from the same patient.

### Calculation of ECM and YAP scores

For ECM scores, the median patient-level expression of stromal Fibronectin, Collagen-I, SMA, Laminin and Vinculin was determined across the full patient dataset. Next, each patient was assigned one point for each instance that the expression of each of the above markers was above the dataset median. The sum of all points was determined as that patient’s ECM score. YAP scores were determined in the same way, with patients being assigned into the “YAP-High” group if their mean nuclear YAP expression in the tumor epithelium fell above the dataset median. All other patients were assigned into the “YAP-Low” group.

### Survival analysis

Kaplan-Meier analysis was used to compare survival outcomes between patient groups with different phenotypic signatures, with Log-rank test used to measure statistical significance. P ≤ 0.05 was used as a cut-off for statistical significance.

### Quantification and statistical analysis

GraphPad Prism (version 9.3.1) was used for all statistical analyses. Outliers were identified with 0.1 % ROTS and distribution was determined with D’Agostino-Pearson normality test. Two-sample testing was performed using Student’s t-test (unpaired, two-tailed) with Welch’s correction (normally distributed data) or nonparametric Mann–Whitney U-test (non-normally distributed data). Multiple comparisons were performed using ANOVA with Holm-Sidak’s post hoc test (normally distributed data) or Dunnett’s post hoc test (non-normally distributed data). Data are presented as column graphs or dot plots (mean±s.d.). P-values less than 0.05 were considered to be statistically significant.

### Data and material availability

Data supporting the findings of this study are available within the paper and its supplementary information files.

## Acknowledgements

We thank J. Siivonen and P. Laasola for technical assistance and the Ivaska lab for scientific discussion. For services, instrumentation and expertise, we would like to thank the Cell Imaging and Cytometry Core (Turku Bioscience Centre, University of Turku) supported by Biocenter Finland, the Euro-BioImaging Finnish Node (Turku Finland), The Finnish Functional Genomics Centre supported by University of Turku, Åbo Akademi University and Biocenter Finland, The Medical Bioinformatics Centre of Turku Bioscience Centre supported by University of Turku, Abo Akademi University, Biocenter Finland and Elixir-Finland, for the sequencing data analysis. FIMM Digital Microscopy and Molecular Pathology Unit supported by HiLIFE and Biocenter Finland for multiplex fluorescence immunohistochemistry and high-content imaging services. This study has been supported by Molecular Regulatory Networks of Life (R’Life) (330033 JI and SW), Finnish Cancer Institute (K. Albin Johansson Professorship, J.I.); a Research Council of Finland Centre of Excellence program (# 346131, J.I. and S.W.); the Cancer Foundation Finland (J.I.); the Sigrid Juselius Foundation (J.I.); the Research Council of Finland’s Flagship InFLAMES (# 337530 & 357910) and the Jane and Aatos Erkko Foundation (J.I.). JK is supported by the University of Turku Doctoral Program for Molecular Medicine and the Finnish Cultural Foundation. MRC was supported by a Research Council of Finland postdoctoral research grant (# 343239). JRWC. was supported by the European Union’s Horizon 2020 research and innovation programme under the Marie Sklodowska-Curie grant agreement [841973] and an Academy of Finland postdoctoral research grant (338585). HA is supported by a fellowship from Fondazione Umberto Veronesi. GF was supported by a Research Council of Finland postdoctoral research grant (332402) and a Turku Collegium for Science Medicine and Technologies postdoctoral fellowship.GS is supported by ERC-Synergy (Grant# 101071470), AIRC-IG (Grant#22821), AIRC 5x1000 (#22759), the Italian Ministry of University and Research (PRIN202223GSCIT_01/G53D23002570006/20229RM8A_001; COMBINE/G53D23007040001/P2022RH4HH002; PNRR_CN3RNA_SPOKE/G43C22001320007. YAM is supported by the Intramural Research Program of the NIH, National Institute of Diabetes and Digestive and Kidney Diseases (NIDDK).

## Author contributions

Conceptualization: JK, SW, JI. Methodology: JK, JI, RC, KP, SW. Formal Analysis: JK, KP, MRC, YAM, FB, HA, FK, JF, JRWC, GF. Investigation: JK, KP, YAM, HA, JH, KV, EP, MN. Visualization: JK, KP, HA, FK, HH. Resource: HI, SV, AM. Writing: JK, GC, RC, JI. Supervision: AM, HI, SV, GS, RC, SW, JI. Funding: JI.

