## Supplementary data figures for "Restoring mechanophenotype reverts malignant properties of ECM-enriched vocal fold cancer"

Extended Data Figures and Figure Legends:

Extended Data Figure 1, Related to Figure 1.

Extended Data Figure 2, Related to Figure 2.

Extended Data Figure 3, Related to Figure 3.

Extended Data Figure 4, Related to Figure 4.

Extended Data Figure 5, Related to Figure 5.

Extended Data Figure 6, Related to Figure 6

Extended Data Figure 7, Related to Figure 7

**Extended data videos:**

**Video 1**: T1 VFC cell proliferation (single cells) on 0.5 kPa Collagen-Fibronectin coated hydrogel. Imaged using Incucyte (ZOOM) every 2 hours for 116 hours, 10x magnification.

**Video 2**: T3 VFC cell proliferation (single cells) on 0.5 kPa Collagen-Fibronectin coated hydrogel. Imaged using Incucyte (ZOOM) every 2 hours for 116 hours, 10x magnification.

**Video 3**: T1 VFC cell proliferation (single cells) on 50 kPa Collagen-Fibronectin coated hydrogel. Imaged using Incucyte (ZOOM) every 2 hours for 116 hours, 10x magnification.

**Video 4**: T3 VFC cell proliferation (single cells) on 50 kPa Collagen-Fibronectin coated hydrogel. Imaged using Incucyte (ZOOM) every 2 hours for 116 hours, 10x magnification.

**Video 5**: T1 VFC cell proliferation (single cells) on Collagen-Fibronectin coated plastic. Imaged using Incucyte (ZOOM) every 2 hours for 116 hours, 10x magnification.

**Video 6**: T3 VFC cell proliferation (single cells) on Collagen-Fibronectin coated plastic. Imaged using Incucyte (ZOOM) every 2 hours for 116 hours, 10x magnification.

**Video 7**: T1 VFC cell proliferation (single cells) on 0.5 kPa Matrigel coated hydrogel. Imaged using Incucyte (ZOOM) every 2 hours for 116 hours, 10x magnification.

**Video 8**: T3 VFC cell proliferation (single cells) on 0.5 kPa Matrigel coated hydrogel. Imaged using Incucyte (ZOOM) every 2 hours for 116 hours, 10x magnification.

**Video 9**: T1 VFC cell proliferation (single cells) on 50 kPa Matrigel coated hydrogel. Imaged using Incucyte (ZOOM) every 2 hours for 116 hours, 10x magnification.

**Video 10**: T3 VFC cell proliferation (single cells) on 50 kPa Matrigel coated hydrogel. Imaged using Incucyte (ZOOM) every 2 hours for 116 hours, 10x magnification.

**Video 11**: T1 VFC cell proliferation (single cells) on Matrigel coated plastic. Imaged using Incucyte (ZOOM) every 2 hours 116 hours, 10x magnification.

**Video 12**: T3 VFC cell proliferation (single cells) on Matrigel coated plastic. Imaged using Incucyte (ZOOM) every 2 hours for 116 hours, 10x magnification.

**Video 13**: NC cell proliferation (colony) on Collagen-Fibronectin coated plastic. Growth medium supplemented with anti-α3α6 antibody (P1B5 and P5G10, 10 µg/ml) at 17h. Imaged using Incucyte (S3) every 60 minutes for 23 hours, 20x magnification.

**Video 14**: T1 VFC cell proliferation (colony) on Collagen-Fibronectin coated plastic. Growth medium supplemented with anti-α3α6 antibody ((P1B5 and P5G10, 10 µg/ml) at 17h. Imaged using Incucyte (S3) every 60 minutes for 23 hours, 20x magnification.

**Video 15**: T3 VFC cell proliferation (colony) on Collagen-Fibronectin coated plastic. Growth medium supplemented with anti-α3α6 antibody (P1B5 and P5G10, 10 µg/ml) at 17h. Imaged using Incucyte (S3) every 60 minutes for 23 hours, 20x magnification.

**Video 16**: NC cell proliferation (colony) on Collagen-Fibronectin coated plastic. Growth medium supplemented with anti-E-cadherin antibody (DECMA-1) at 17h. Imaged using Incucyte (S3) every 60 minutes for 23 hours, 20x magnification.

**Video 17**: T1 VFC cell proliferation (colony) on Collagen-Fibronectin coated plastic. Growth medium supplemented with anti-E-cadherin antibody (DECMA-1) at 17h. Imaged using Incucyte (S3) every 60 minutes for 23 hours, 20x magnification.

**Video 18**: T3 VFC cell proliferation (colony) on Collagen-Fibronectin coated plastic. Growth medium supplemented with anti-E-cadherin antibody (DECMA-1) at 17h. Imaged using Incucyte (S3) every 60 minutes for 23 hours, 20x magnification.

**Video 19**: NC cell proliferation (monolayer) on Collagen-Fibronectin coated plastic. Growth medium supplemented with anti-IgG antibody (10 µg/ml). Imaged using Incucyte (S3) every 10 minutes for 24 hours, 20x magnification.

**Video 20:** NC cell proliferation (monolayer) on Collagen-Fibronectin coated plastic. Growth medium supplemented with anti-α3 integrin antibody (P1B5, 10 µg/ml). Imaged using Incucyte (S3) every 10 minutes for 24 hours, 20x magnification.

**Video 21:** NC cell proliferation (monolayer) on Collagen-Fibronectin coated plastic. Growth medium supplemented with anti-α6 integrin antibody (P5G10, 10 µg/ml). Imaged using Incucyte (S3) every 10 minutes for 24hours, 20x magnification.

**Video 22:** NC cell proliferation (monolayer) on Collagen-Fibronectin coated plastic. Growth medium supplemented with anti-α3α6 integrin antibody (P1B5 and P5G10, 10 µg/ml). Imaged using Incucyte (S3) every 10 minutes for 24 hours, 20x magnification.

**Video 23:** NC cell proliferation (monolayer) on Collagen-Fibronectin coated plastic. Growth medium supplemented with anti-β1 integrin antibody (mAb13, 10 µg/ml). Imaged using Incucyte (S3) every 10 minutes for 24 hours, 20x magnification.

**Video 24:** T1 VFC cell proliferation (monolayer) on Collagen-Fibronectin coated plastic. Growth medium supplemented with anti-IgG antibody (10 µg/ml). Imaged using Incucyte (S3) every 10 minutes for 24 hours, 20x magnification.

**Video 25:** T1 VFC cell proliferation (monolayer) on Collagen-Fibronectin coated plastic. Growth medium supplemented with anti-α3 integrin antibody (P1B5, 10 µg/ml). Imaged using Incucyte (S3) every 10 minutes for 24 hours, 20x magnification.

**Video 26:** T1 VFC cell proliferation (monolayer) on Collagen-Fibronectin coated plastic. Growth medium supplemented with anti-α6 integrin antibody (P5G10, 10 µg/ml). Imaged using Incucyte (S3) every 10 minutes for 24 hours, 20x magnification.

**Video 27:** T1 VFC cell proliferation (monolayer) on Collagen-Fibronectin coated plastic. Growth medium supplemented with anti-α3α6 integrin antibody (P1B5 and P5G10, 10 µg/ml). Imaged using Incucyte (S3) every 10 minutes for 24 hours, 20x magnification.

**Video 28:** T1 VFC cell proliferation (monolayer) on Collagen-Fibronectin coated plastic. Growth medium supplemented with anti-β1 integrin antibody (mAb13, 10 µg/ml). Imaged using Incucyte (S3) every 10 minutes for 24 hours, 20x magnification.

**Video 29:** T3 VFC cell proliferation (monolayer) on Collagen-Fibronectin coated plastic. Growth medium supplemented with anti-IgG antibody (10 µg/ml). Imaged using Incucyte (S3) every 10 minutes for 24 hours, 20x magnification.

**Video 30:** T3 VFC cell proliferation (monolayer) on Collagen-Fibronectin coated plastic. Growth medium supplemented with anti-α3 integrin antibody (P1B5, 10 µg/ml). Imaged using Incucyte (S3) every 10 minutes for 24 hours, 20x magnification.

**Video 31:** T3 VFC cell proliferation (monolayer) on Collagen-Fibronectin coated plastic. Growth medium supplemented with anti-α6 integrin antibody (P5G10, 10 µg/ml). Imaged using Incucyte (S3) every 10 minutes for 24 hours, 20x magnification.

**Video 32:** T3 VFC cell proliferation (monolayer) on Collagen-Fibronectin coated plastic. Growth medium supplemented with anti-α3α6 integrin antibody (P1B5 and P5G10, 10 µg/ml). Imaged using Incucyte (S3) every 10 minutes for 24 hours, 20x magnification.

**Video 33:** T3 VFC cell proliferation (monolayer) on Collagen-Fibronectin coated plastic. Growth medium supplemented with anti-β1 integrin antibody (mAb13, 10 µg/ml). Imaged using Incucyte (S3) every 10 minutes for 24 hours, 20x magnification.

### Extended data Figures

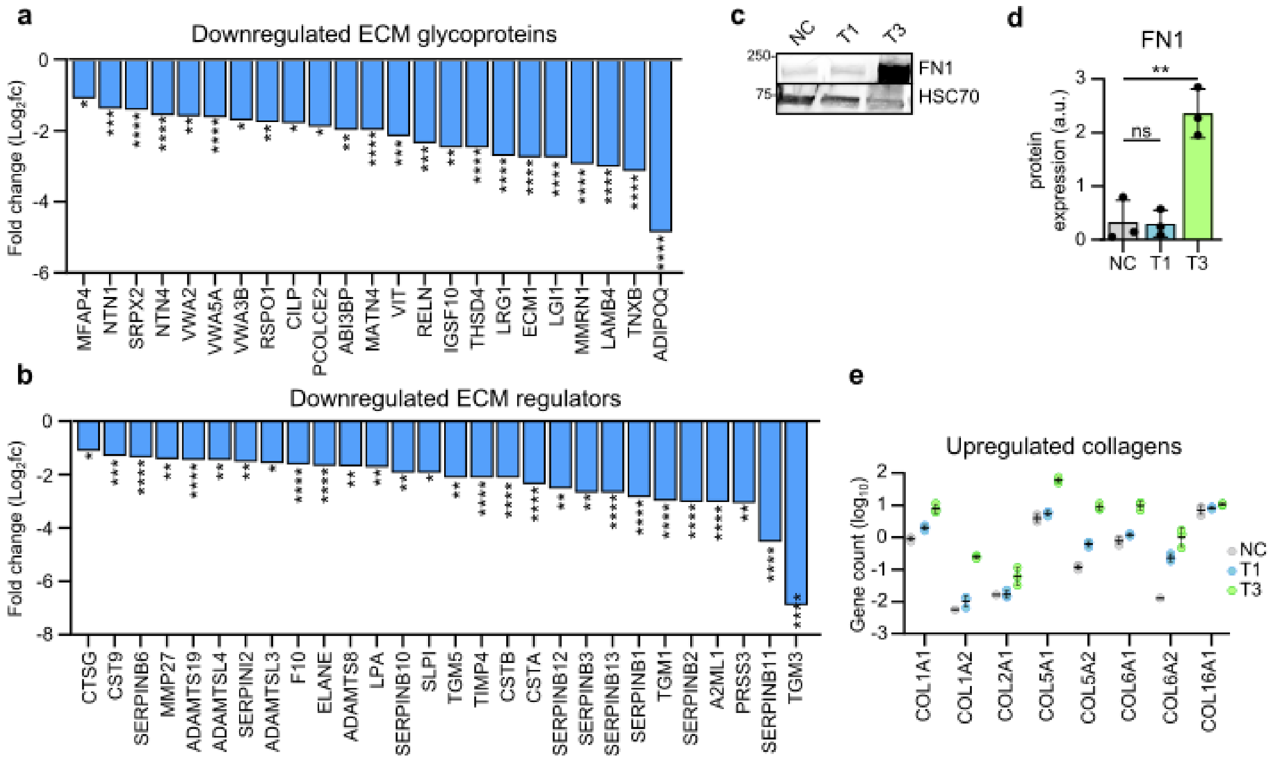


**Extended Data Figure 1 | Vocal fold cancer is associated with elevated gene expression of ECM components and stiffening of tissue**

**a & b** Differentially downregulated (fold change, log2) ECM glycoproteins (**a**) and ECM regulators (**b**) in VFC tissue (T1-T4, n=54) compared to normal tissue (n=12) in patients (TCGA-data, FDR < 0.05). **c & d**, Representative immunoblot (**c**) and quantification (**d**) of fibronectin protein expression (mean±s.d., n=3) in NC cells and vocal fold T1 and T3 cancer cells. **e**, Upregulated gene count (log10, RNA-seq) of collagens in NC cells and vocal fold T1 and T3 cancer cells (n=3). Data are mean (± s.d.). FDR was used to asses statistical significance of differentially expressed genes and ordinary one-way Anova followed by post hoc Dunnett's multiple comparisons test was used to asses statistical significance of protein expression.

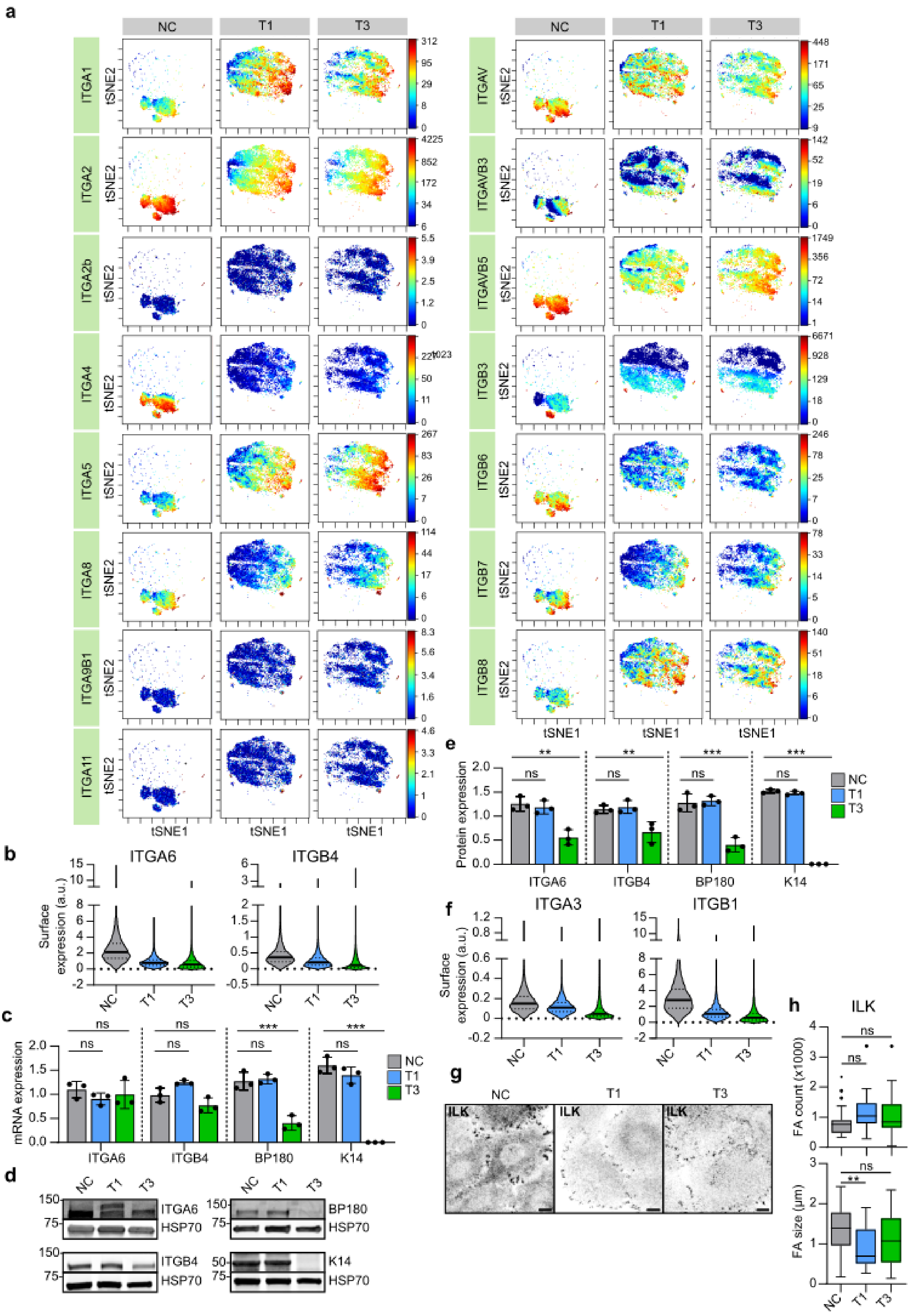
a
NC
T1
T3
ITGA1
ITGA2
ITGA2b
ITGA4
ITGA5
ITGA8
ITGA9B1
ITGA11
ITGAV
ITGAVB3
ITGAVB5
ITGB3
ITGB6
ITGB7
ITGB8
tSNE2
tSNE1
b
ITGA6
ITGB4
Surface expression (a.u.)
c
mRNA expression
ITGA6
ITGB4
BP180
K14
d
ITGA6
HSP70
ITGB4
BP180
K14
e
Protein expression
f
ITGA3
ITGB1
g
ILK
h
ILK
FA count (x1000)
FA size (µm)
ns
**
***

**Extended Data Figure 2 | Expression and subcellular localization of laminin-binding integrins is altered in vocal fold cancer**

**a**, t-SNE visualization of ITGA1, ITGA2, ITGA2b, ITGA4, ITGA5, ITGA8, ITGA9B1, ITGA11, ITGAV, ITGAVB3, ITGAVB5, ITGB3, ITGB6, ITGB7 and ITGB8 single-cell surface expression (MassCytof) in NC cells and vocal fold T1 and T3 cancer cells. **b**, Violin plot representation of mean ITGA6 and ITGB4 single-cell surface expression (MassCytof) in NC cells and vocal fold T1 and T3 cancer cells. **c**, Relative ITGA6, ITGB4, BP180 and K14 mRNA expression levels (gene count) in NC cells and vocal fold T1 and T3 cancer cells (n=3). **d & e**, Representative immunoblots (**d**) and quantification (**e**) of relative ITGA6, ITGB4, BP180 and K14 protein expression levels in NC cells and vocal fold T1 and T3 cancer cells (n=3). **f**, Violin plot representation of mean ITGA3 and ITGB1 single-cell surface expression (MassCytof) in NC cells and vocal fold T1 and T3 cancer cells. **g**, Representative ILK confocal immunofluorescence images of NC cells and VFC T1 and T3 cells (n=3). Scale bar 10 µm. **h**, Quantification of FA number (count) (left) and size (right) using ILK as a marker in NC cells (n=28), and VFC T1 (n=29-30) and T3 (n=28-30) cells. Data are mean box plots (± s.d.) or tukey box plots**.** n is the total number of average FA count/size per cell in FOV pooled from three independent experiments Statistical significance was assessed using Kruskal-Wallis test followed by post hoc Dunn's multiple comparisons test.

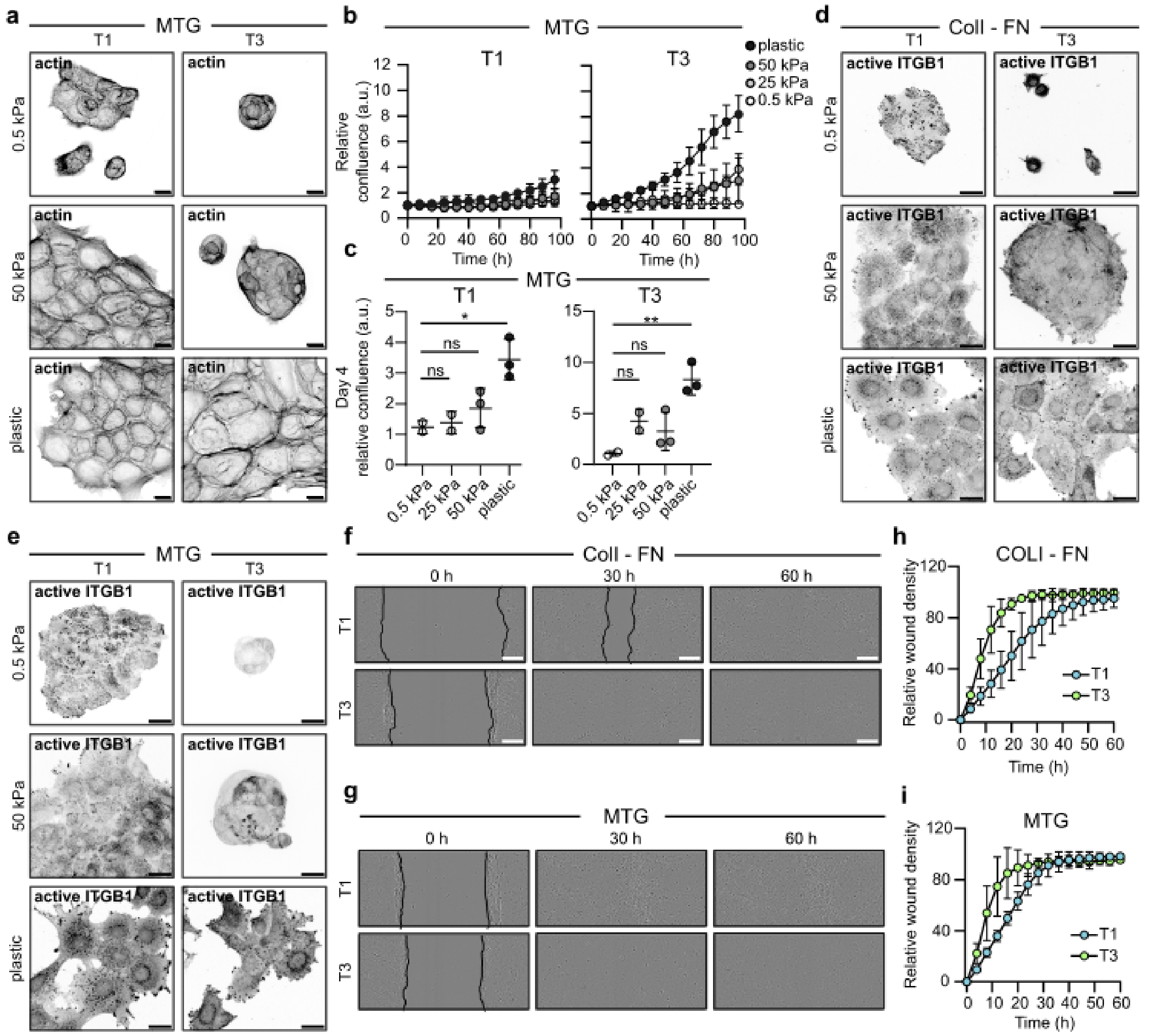


**Extended Data Figure 3 | Stiffening of vocal fold tissue supports increased cell proliferation, migration and invasion**

**a,** Representative actin confocal immunofluorescence images of T1 and T3 VFC cells on 0.5 and 50 kPa hydrogels and plastic coated with Matrigel (n=3). Scale bar 50 µm. **b & c**, Proliferation (**b**) of T1 and T3 VFC cells on hydrogels of varying stiffnesses (0.5 kPa, 25 kPa, 50 kPa) and plastic and confluence at end-point (**c**). **d & e**, Representative active ITGB1 (12G10) confocal immunofluorescence images of T1 and T3 VFC cells on 50 kPa hydrogels coated with collagen and fibronectin (**d**) or Matrigel (**e**) (n=3). Scale bar 50 µm. **f & g**, Representative phase-contrast images of T1 and T3 VFC cells wound healing assay at different timepoints (0h, 30h and 60h) on collagen I and fibronectin (**f**) or Matrigel (**g**) coated plates. Scale bar 100 µm. (n=3). **h & i**, Relative wound density of T1 and T3 VFC cells on collagen I and fibronectin (**h**) or Matrigel (**i**) coated plates. (n=3). Data are mean (± s.d.). Statistical significance was assessed using Kruskal-Wallis test followed by post hoc Dunn's multiple comparisons test.

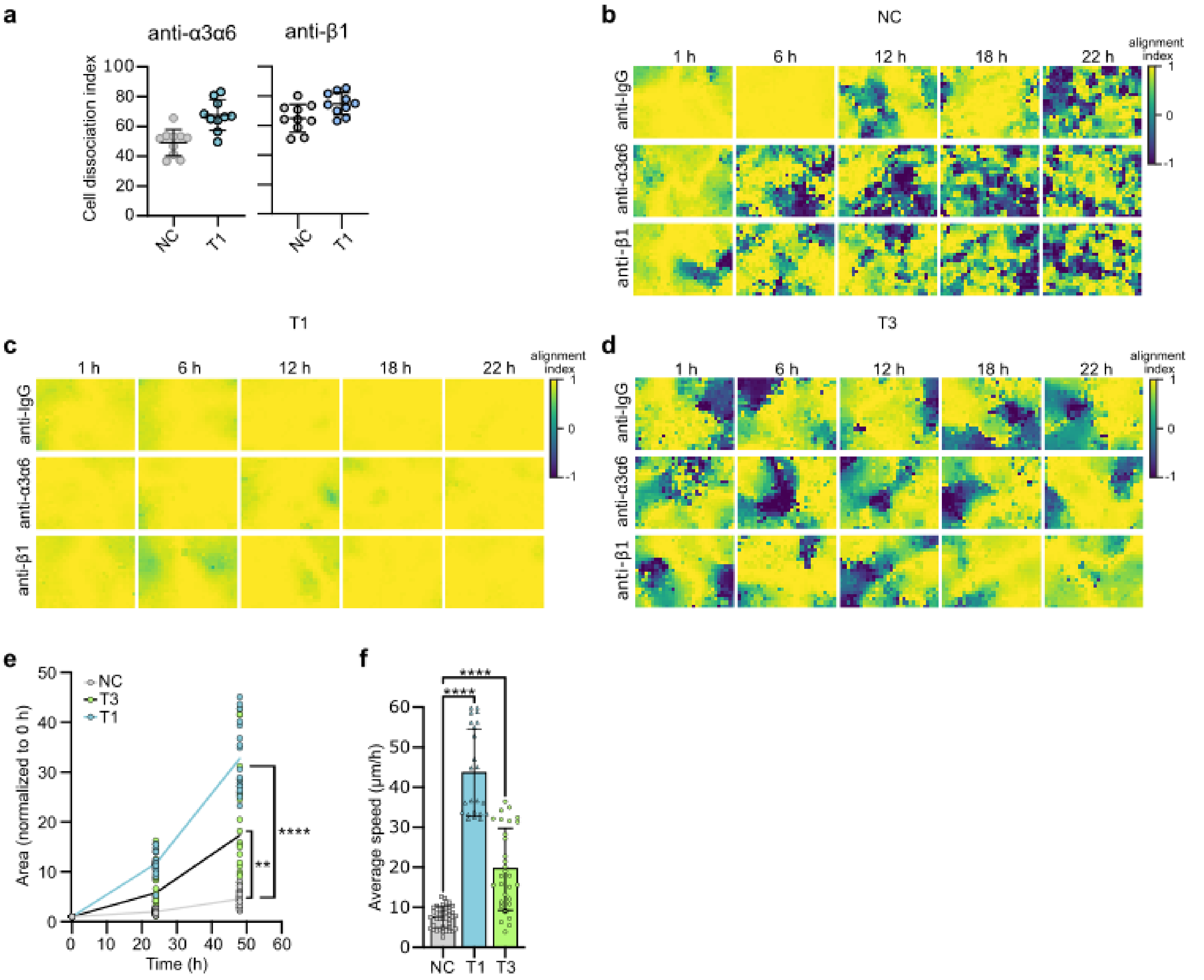


**Extended Data Figure 4 | Inhibition of laminin-binding integrins modulates monolayer dynamics and disrupts cell clustering in 3D-spheroids**

**a** Quantification of cell dissociation index in NC and VFC T1 cells treated with anti-α3α6 or anti-β1 integrin blocking antibodies (n=3). **b-d**, Graphic visualization of cell velocity alignment index in NC (**b**), VFC T1 (**c**) and T3 (**d**) cells treated with anti-α3α6 or anti-β1 integrin blocking antibodies. **e & f**, Quantification of normalized wetting area (**e**) and average wetting speed (μm/h) (**f**) of NC and VFC T1 and T3 cells (n=3). Data are mean ± s.d. Statistical significance was assessed using Kruskal-Wallis test followed by post hoc Dunn's multiple comparisons test.

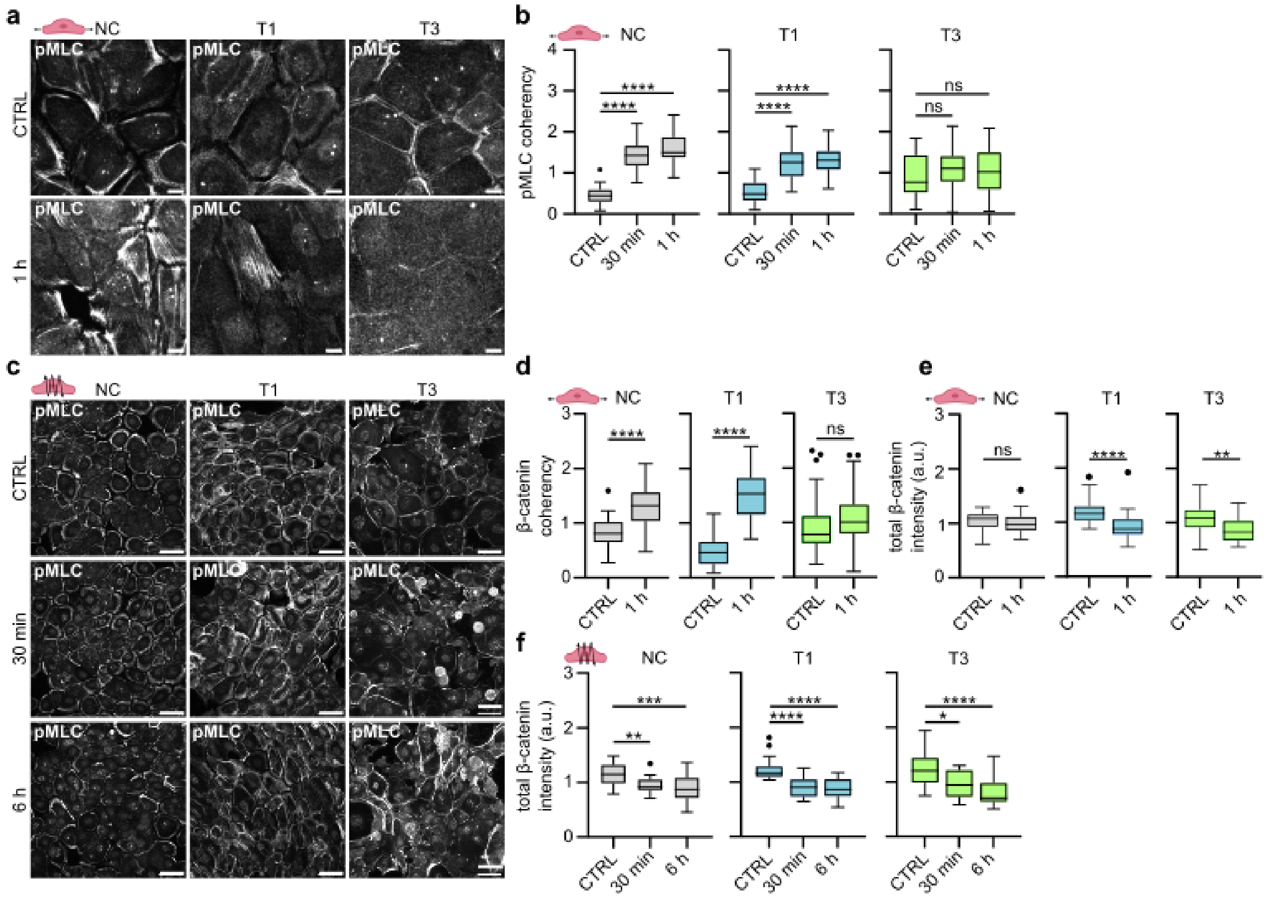


**Extended Data Figure 5 | Mechanical stimuli induce cytoskeletal and junctional alterations and cell extrusion in VFC**

**a**, Representative pMLC confocal immunofluorescence images of NC cells and VFC T1 and T3 cells subjected to stretch (n=3). **b**, Quantification of pMLC coherency of NC cells and vocal fold T1 and T3 cells subjected to stretching (n=3). **c**, Representative pMLC confocal immunofluorescence images of NC cells and VFC T1 and T3 cells subjected to vibration (n=3). Scale bar 20 µm. **d**, Quantification of β-catenin coherency of NC cells and VFC T1 and T3 cells subjected to stretching (n=3). **e & f**, Quantification of total β-catenin intensity of NC cells and VFC T1 and T3 cells subjected to stretching (**e**) or vibration (f) (n=3). Data are illustrated as tukey box plots or violin blots (average of 8 FOV's pooled from three independent experiments). Kruskal-Wallis test followed by post hoc Dunn's multiple comparisons test was used to asses statistical significance.

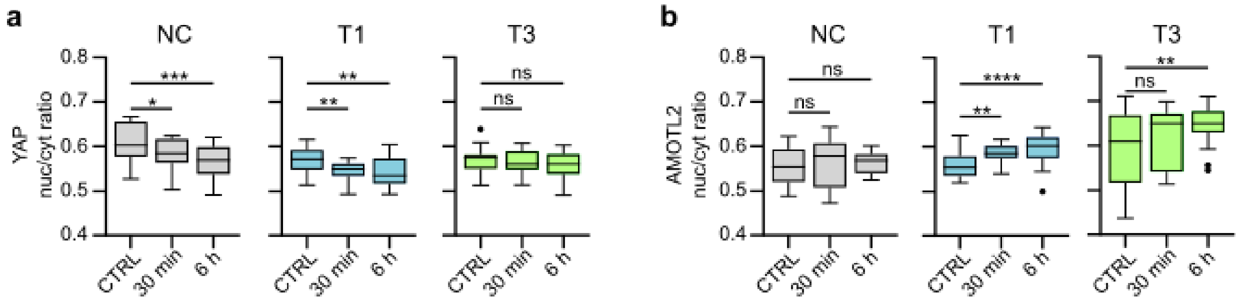


**Extended Data Figure 6 | Phonomimetic mechanical stimuli decreases nuclear and total YAP levels**

**a**, Quantification of YAP nuclear to cytoplasmic ratio in NC cells and VFC T1 and T3 cells subjected to vibration (50-250 Hz, 1 min on/off) for 30 min or 6h compared to non-vibrated control. **b**, Quantification of AMOTL2 nuclear to cytoplasmic ratio in NC cells and VFC T1 and T3 cells subjected to vibration (50-250 Hz, 1 min on/off) for 30 min or 6h compared to non-vibrated control. Data are illustrated as tukey box plots (average of 8 FOV’s pooled from three independent experiments). Ordinary one-way Anova followed by post hoc Dunnett's multiple comparisons test was used to asses statistical significance.

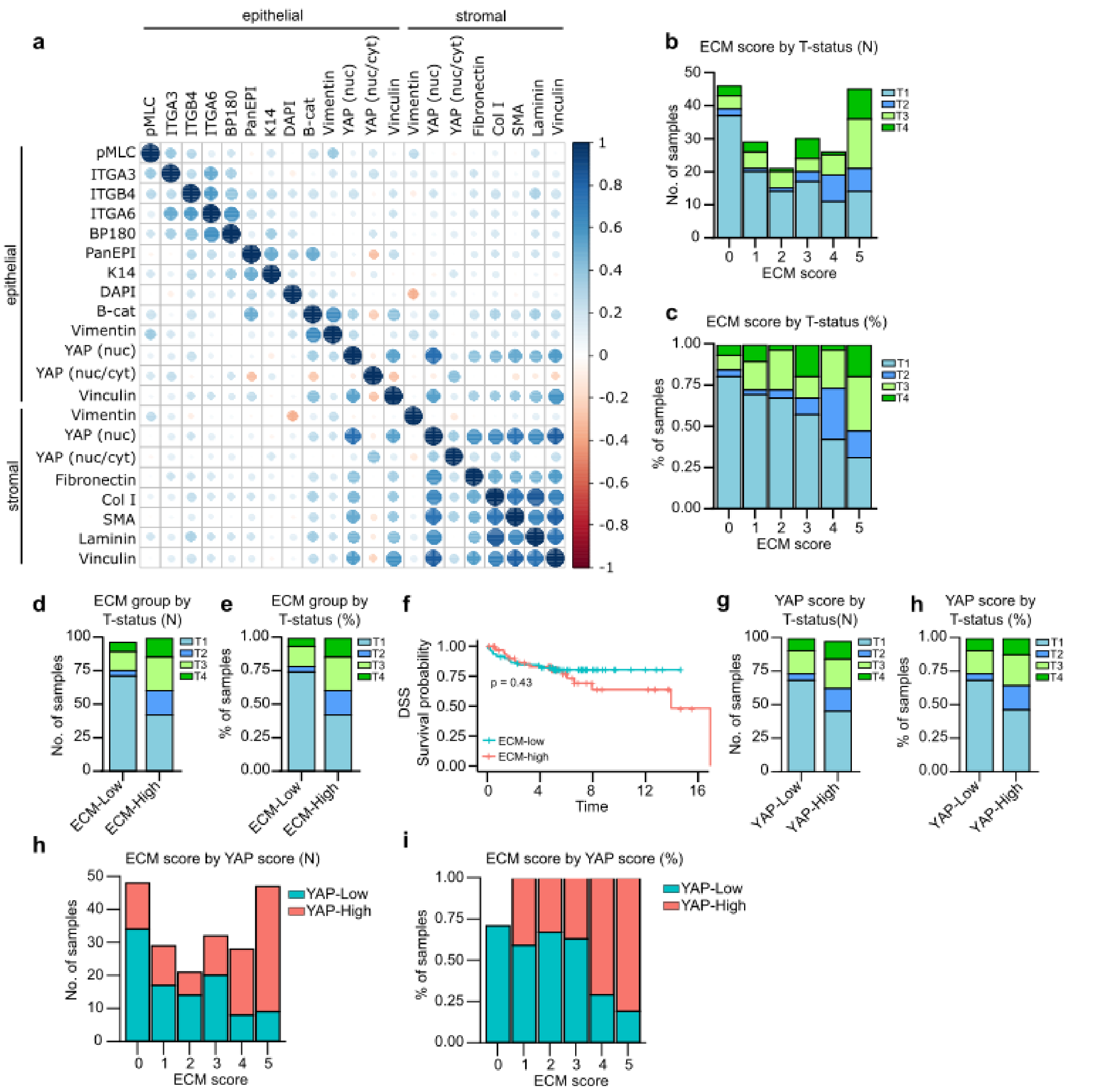


**Extended Data Figure 7 | High YAP levels correlate with high ECM expression and poor disease specific survival**

**a**, Patient-level correlation of epithelial (pMLC, ITGA3, ITGB4, ITGA6, BP180, PanEpi, K14, dapi, vimentin, YAP and vinculin) and stromal (vimentin, YAP, fibronectin, COL I, SMA, laminin and vinculin) marker mean expression in TMA multiplex histology. **b**, ECM-score by T-status illustrated as number of samples (N). **c**, ECM score by T-statusillustrated as percentage of samples (%). **d**, ECM group by T-status illustrated as number of samples (N). **e**, ECM group (ECM-low and ECM-high) by T-status illustrated as percentage of samples (%). **f**, Disease specific survival of ECM-high and ECM-low patients.  **g & h**, YAP score by T-status illustrated as number of samples (N) (**g**) and percentage of samples (%) (**h**). **i & j,** ECM score by YAP score illustrated as number of samples (N) (**i**) and percentage of samples (%) (**j**). Data are mean expression and statistical significance of Kaplan-Meier analysis was assessed with Log-rank test.
